# DPLM: Dynamics-aware Protein Language Model via Contrastive Learning between Sequence and Molecular Dynamics Simulation Trajectory

**DOI:** 10.64898/2026.04.29.721692

**Authors:** Yuexu Jiang, Duolin Wang, Newton A. Ihoeghian, Ibrahim A. Imam, Jin Chen, Dong Xu, Qing Shao

## Abstract

Protein dynamics captures the interactions between residues and the conformational changes they undergo. Encoding such information could enhance a deep learning model’s ability to predict protein properties. Here we introduce a dynamics-aware protein language model (DPLM) that encodes protein dynamics into a sequence-derived representation. We represent each molecular dynamics (MD) trajectory as a sequence of backbone Cα contact maps and encode it with a Video Vision Transformer (ViViT). We then align the ViViT-derived dynamics embedding with the ESM2-derived sequence embedding using dual-view contrastive learning. The root-mean-square fluctuation correlation and dynamic cross-correlation matrix reconstruction confirm that contrastive learning incorporates dynamics information into the sequence embedding. We applied DPLM as a protein representation backbone for predicting diverse dynamics-related downstream tasks, including liquid-liquid phase separation, cryptic-site detection, and mutation effect on protein thermal stability. DPLM shows strong performance, especially where the evolutionary or structural signal is weak. These results show that learning the geometry of molecular motion yields transferable dynamic information that complements the evolutionary and structural features of existing PLMs.

**Significance Statement:** Proteins perform their functions in constant motion. Most protein language models (PLMs) miss this dynamic dimension, learning only from sequence or fixed structure. A few emerging PLMs model dynamics through conformational snapshots or derived dynamic properties rather than the full time-resolved motion. As a result, the temporal ordering inherent in molecular motion is not explicitly captured. We introduce DPLM, a dynamics-aware protein language model that uses self-supervised learning to align protein sequences with their molecular dynamics simulation trajectories. The resulting sequence-based representation encodes general dynamic information and benefits diverse downstream tasks, especially where evolution and structure alone fall short. DPLM offers a general route to physics-informed protein representation.

## Introduction

Proteins carry out most molecular functions in the cell^1^. Predicting protein properties is a central problem in computational biology^2^. Protein language models (PLMs) have emerged as powerful tools for understanding and designing proteins^3^. PLMs adapt large-scale self-supervised learning from natural language processing to capture statistical and functional regularities encoded in protein sequences^4–7^. The resulting contextual representations transfer effectively across diverse downstream tasks^8,9^. A growing class of PLMs align sequences with structures^10–13^ or Enzyme Commission (EC) number annotations^14^ to make them structure-aware or EC-aware. Despite this success, mainstream PLMs are still trained on static descriptions of proteins. But proteins are dynamic molecules, and nearly 30% of the human proteome consists of intrinsically disordered regions lacking static structures^15^. PLMs that ignore protein dynamics therefore lack information central to properties such as conformational flexibility, binding affinity, condensate formation and thermodynamic stability^16–18^. Molecular dynamics (MD) simulations model protein motion explicitly, resolving residue-level flexibility and collective motions that sequence or static structure cannot reveal^19^. But MD is computationally expensive, especially for large systems and long simulations. The field therefore needs a way to incorporate dynamics into sequence-based representations without running an MD simulation for every protein.

A few recent efforts have started to move in this direction. SeqDance and ESMDance^20^ use supervised objectives to regress 23 predefined dynamic properties, including Root-Mean-Square Fluctuation (RMSF), Cα co-movement correlations and interaction-frequency profiles, extracted from MD or normal-mode analysis of roughly 64,000 proteins. ESMDance builds these targets on top of ESM2 outputs and clearly outperforms the SeqDance variant trained from random initialization. Kalifa et al. instead conditioned the protein sequence representation on a generated conformational ensemble^21^. A generative model first proposes multiple plausible conformations for a sequence, and a dynamic embedding layer then integrates over them to yield a single sequence of conformation-aware token embeddings. These methods model protein dynamics through ensembles of static conformations rather than time-resolved physical trajectories; consequently, the temporal ordering and sequential information inherent in molecular motion are not explicitly encoded^22^.

Here we introduce the Dynamics-aware Protein Language Model (DPLM), which embeds protein dynamics into ESM2 sequence-derived representations. DPLM has three key features. First, it uses a Video Vision Transformer (ViViT)^23^ to capture spatial and temporal patterns directly from MD trajectories rather than predefined dynamic properties or static conformational ensembles. Second, DPLM preserves evolutionary information by freezing the ESM2 sequence encoder and injects dynamics through parameter-efficient adapters^24^. Third, it uses dual-view contrastive learning^25^ to align sequence and MD embeddings. This self-supervised training is designed to learn general-purpose representations rather than task-specific ones. The resulting representations require only protein sequences at inference.

We benchmarked DPLM against one representative PLM from each paradigm: the evolution-based ESM2^6^, the structure-aware ProstT5^13^, and the dynamics-supervised ESMDance^20^. All three need only sequence at inference. This design isolates the contribution of dynamics in DPLM, since each baseline is the leading model of an alternative information source. We also compared against task-specific methods where available. DPLM embeddings correlate with RMSF and Dynamic Cross-Correlation Matrix (DCCM) significantly more strongly than the other PLMs. This dynamic signal translates into consistent downstream gains. DPLM achieves the best performance on liquid-liquid phase separation (LLPS) prediction. DPLM complements the structure-aware ProstT5 on cryptic binding-site detection, where both dynamics and structure matter. DPLM also predicts mutation-induced changes in thermal stability for proteins with limited evolutionary information better than other PLMs. A pattern is clear that the advantage of DPLM is largest where evolutionary and structural signals are weakest, such as proteins unseen in training sets among tasks, *de novo* designed proteins, and rapidly evolving viral sequences with sparse homology. Together, these results show that DPLM addresses a key limitation of PLMs trained on static descriptions. DPLM could be a powerful protein representation method for predicting dynamics-related properties.

## Results

### An overview of the DPLM framework

We collected the matched protein sequence and MD trajectory data from the Atlas database^26^. Before training, we processed and split the ATLAS dataset into training and test sets with sequence-homology control. Figure 1 shows the DPLM architecture and training process. We cut the protein sequences into fragments and clipped the matched MD trajectories to fit the ViViT input dimensions (see Methods for data and processing details). DPLM couples two encoders in a SimCLR-style contrastive framework^25^. We modified it to address cross-view contrast as introduced in X-CLIP^27^ (Methods). On the sequence side, a frozen ESM2-650M backbone with adapter tuning works as the sequence encoder. It converts the amino-acid sequence into residue-level representations, further pooled into protein-level representations. On the MD side, no pretrained foundation model exists for MD-trajectory embedding. We therefore rendered each trajectory as a “contact-motion video” (a time-ordered stack of residue–residue contact maps) and used ViViT (google/vivit-b-16×2-kinetics400) as the MD encoder to convert the trajectory into a protein-level representation. Each branch is followed by a lightweight projection head. Our dual-view contrastive learning aligns the two projected embeddings. The objective is to pull together the sequence and MD embeddings of the same protein while pushing apart those of different proteins. We also ran a comprehensive, systematic ablation test to evaluate the contribution of each module in the DPLM architecture. The ablation test report is in the Supplementary Materials.

**Fig 1.**
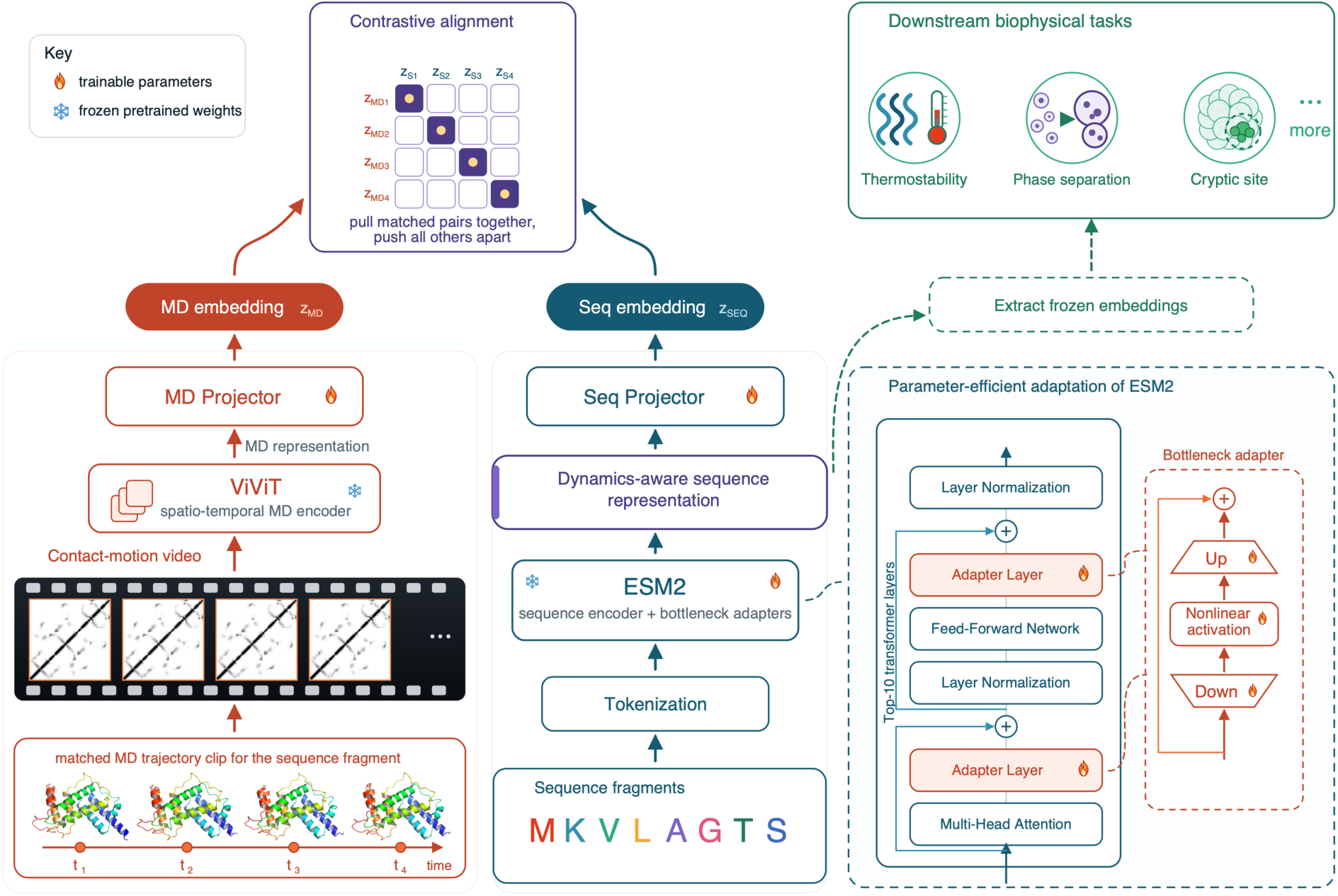
Overview of the dynamics-aware protein language model (DPLM). During training, protein fragments and matched MD trajectory clip are fed into the model. A multi-view contrastive objective aligns the sequence and MD embeddings of the same protein while pushing apart embeddings from different proteins. After pretraining, the MD branch is discarded and the protein representation from the sequence encoder is used directly for downstream tasks (dashed arrow). Flame symbols denote trainable modules, all other backbones are frozen. Adapters are inserted into the top 10 transformer layers of ESM-2 650M (the 10 layers closest to the output in its 33-layer stack).

The alignment between sequence and MD is self-supervised, and the pretrained backbones (ESM2 and ViViT) remain frozen. We trained only the adapters and the projection heads. This parameter-efficient tuning allows DPLM to use a small amount of training data to inject dynamics into the sequence representation. It also reduces the risk of overwriting the evolutionary knowledge in ESM2 and the spatiotemporal relations in ViViT. After pretraining, we extracted the adapted sequence encoder for inference without the need of MD simulation. The adapted sequence encoder (Fig. 1, dashed arrow before the projection layer) provides the dynamics-aware embeddings for downstream tasks, at the residue or protein level as each task requires. We provide the full architectural and training details in the Methods section.

### Dynamic features captured in DPLM representations

To test whether DPLM encodes protein dynamics in its sequence representation, we probed the learned embeddings against MD-derived properties, the residue RMSF and the DCCM. We compared DPLM with ESM2, ProstT5 and ESMDance. Unless otherwise noted, DPLM and ESM2 denote the 650M-parameter models; ProstT5 (3B) and ESMDance (35M) are used at their only available scales.

For each residue, we reduced its embedding to a single residue embedding scalar, defined as the L2 norm of the residue’s embedding vector (Equation (3) in Methods). We calculated the correlation between the embedding scalar and the residue’s RMSF using Spearman’s rank correlation (Equation (4) in Methods). Pooled over all residues in the ATLAS test proteins (Fig. 2a), DPLM correlates with RMSF more strongly (ρ = 0.32) than ESM2 (ρ = 0.23), ProstT5 (ρ = 0.18) or ESMDance (ρ = 0.05). Computed separately within each of the 20 amino-acid types (Fig. 2b), DPLM again ranked highest, significantly ahead of the other methods. Within each protein (Fig. 2c), the gap between DPLM and ESM2 narrowed because most residues are stable, but DPLM stayed significantly above the other methods. We next tested whether the embedding scalar also captures differences in intrinsic flexibility among amino acid types. The per-residue RMSF and DPLM embedding scalar co-vary across the amino-acid series (Fig. 2e). Buried hydrophobic and aromatic residues (Trp, Phe, Tyr, Ile, Val, Leu) are stable and have relatively low embedding scalars. Flexible residues (Gln, Lys, Glu) have the highest embedding scalars.

**Fig 2.**
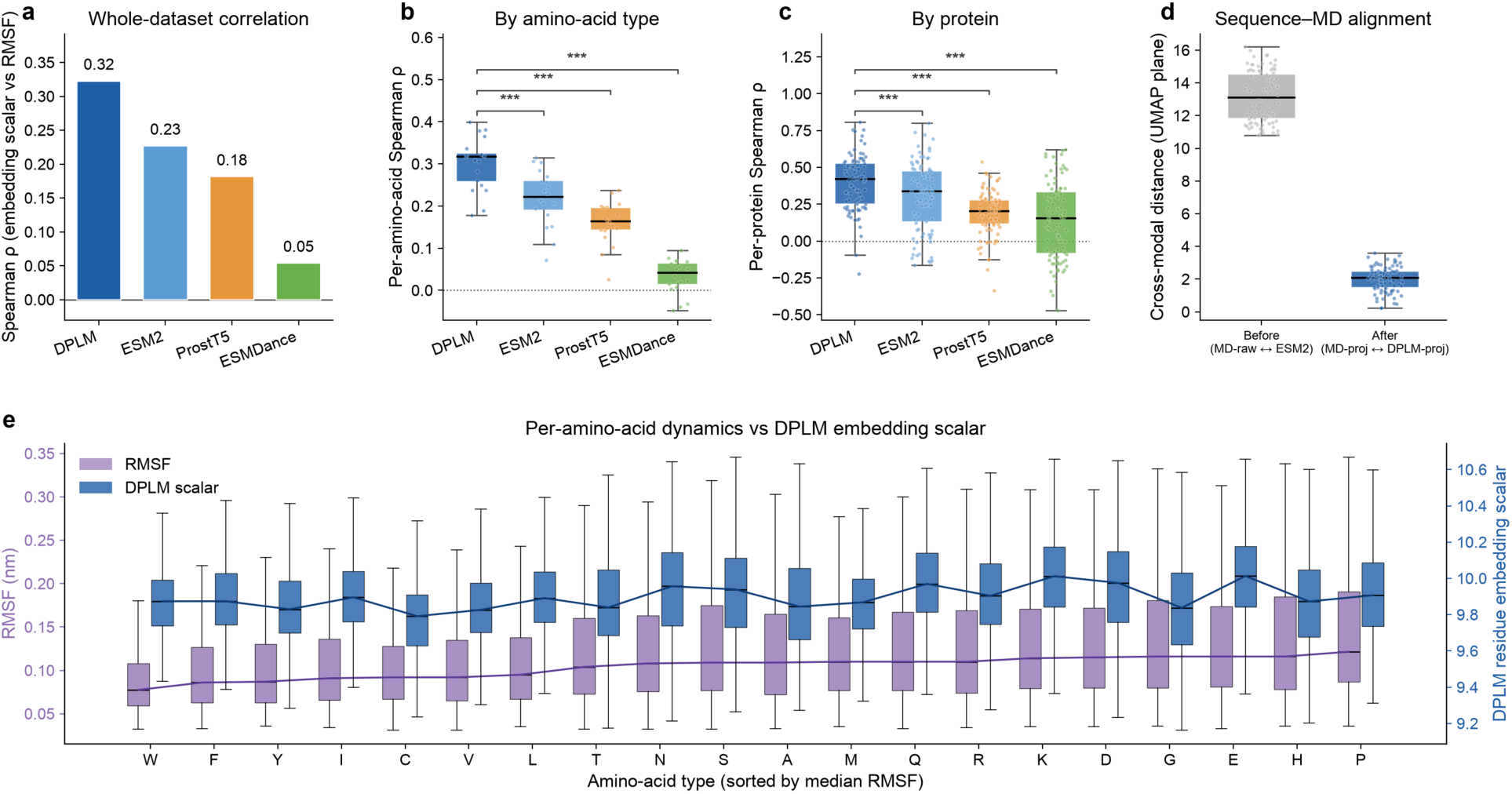
DPLM representation encodes dynamics. **a.** The Spearman ρ between per-residue embedding scalar and RMSF, pooling all Atlas testing dataset residues. **b.** The Spearman ρ computed separately within each of the 20 amino-acid types. Points are individual amino acid types. **c.** The Spearman ρ computed separately within each of the 92 test proteins. Points are individual proteins. **d.** Cross-modal distance between paired sequence and MD embeddings in a joint UMAP plane, before and after the contrastive projection. **e.** For each amino acid, the distribution of per-residue RMSF (purple, left axis) and of the DPLM per-residue embedding scalar (blue, right axis) are shown side by side, with amino acids ordered left to right by increasing median RMSF.

At the protein level, we tested how the contrastive objective affects the geometry of the representation space. We embedded each ATLAS test protein in four ways: the raw MD (ViViT) embedding, the MD embedding after the trained MD projection head, the raw ESM2 sequence embedding, and the sequence embedding after the trained sequence projection head (the DPLM embedding). We then measured the distance between the paired sequence and MD views in a joint cosine-UMAP plane, before and after projection (Fig. 2d). The distance drops significantly after learning. Contrastive pretraining thus reshapes the sequence representation to align with MD information. The alignment is systematic but not exact. Each protein contributes three MD replicates with differing embeddings, so the projected sequence and MD views converge into neighboring regions rather than collapsing onto one another, capturing the shared dynamic signal while preserving replicate-level variation. Supplementary Fig. S3 shows two-dimensional UMAP visualizations of all four views for the three MD replicates. Figure 2 reports one MD replicate; results for all three replicates appear in Supplementary Figs. S1 and S2.

Beyond the flexibility of individual residues, the DCCM captures pairwise co-movement between residues (defined in Equation (5) in Methods). We reconstructed DCCM using representations from different PLMs and correlated it with the ground-truth DCCM computed from the reference MD trajectory. Because the DCCM is defined in terms of temporal correlations, this metric directly probes the benefit of pretraining DPLM on native MD trajectories.

For each held-out test protein, we first correlated the pairwise cosine similarity of residue embeddings with the ground-truth DCCM, without fitting any parameters (Fig. 3a; Methods). The structure-aware ProstT5 was clearly strongest (median Pearson r = 0.28), followed by DPLM (0.12), ESMDance (0.09) and ESM2 (0.08). This ordering is expected as spatially connected residues tend to move together. The ranking reversed when we weighted the same read-out by each PLM’s attention maps, again without training (Fig. 3b; Methods). DPLM has now recovered the DCCM best, well above ProstT5. Its lead over ESM2 and ESMDance is small in distribution but significant by paired Wilcoxon test: DPLM won on 77% of proteins against ESM2, 60% against ESMDance and 100% against ProstT5 (Fig. 3c). Adding attention improved every method, but the ESM-family backbones gained far more (roughly +0.40, versus +0.12 for ProstT5). This asymmetry reflects where each model stores relationship information. ESM2-based models are masked-language-model backbones, in which contact-like couplings emerge in the self-attention maps^28^. ProstT5 is a T5 encoder trained to translate between amino-acid and 3Di structure tokens, so structural information enters through the token vocabulary and is already present in the embeddings. DPLM’s advantage over the PLMs held across the protein-length range, and its margin over ESMDance grew with sequence length (Fig. 3d). This indicates that DPLM can detect long-range amino acid co-movement.

**Fig 3.**
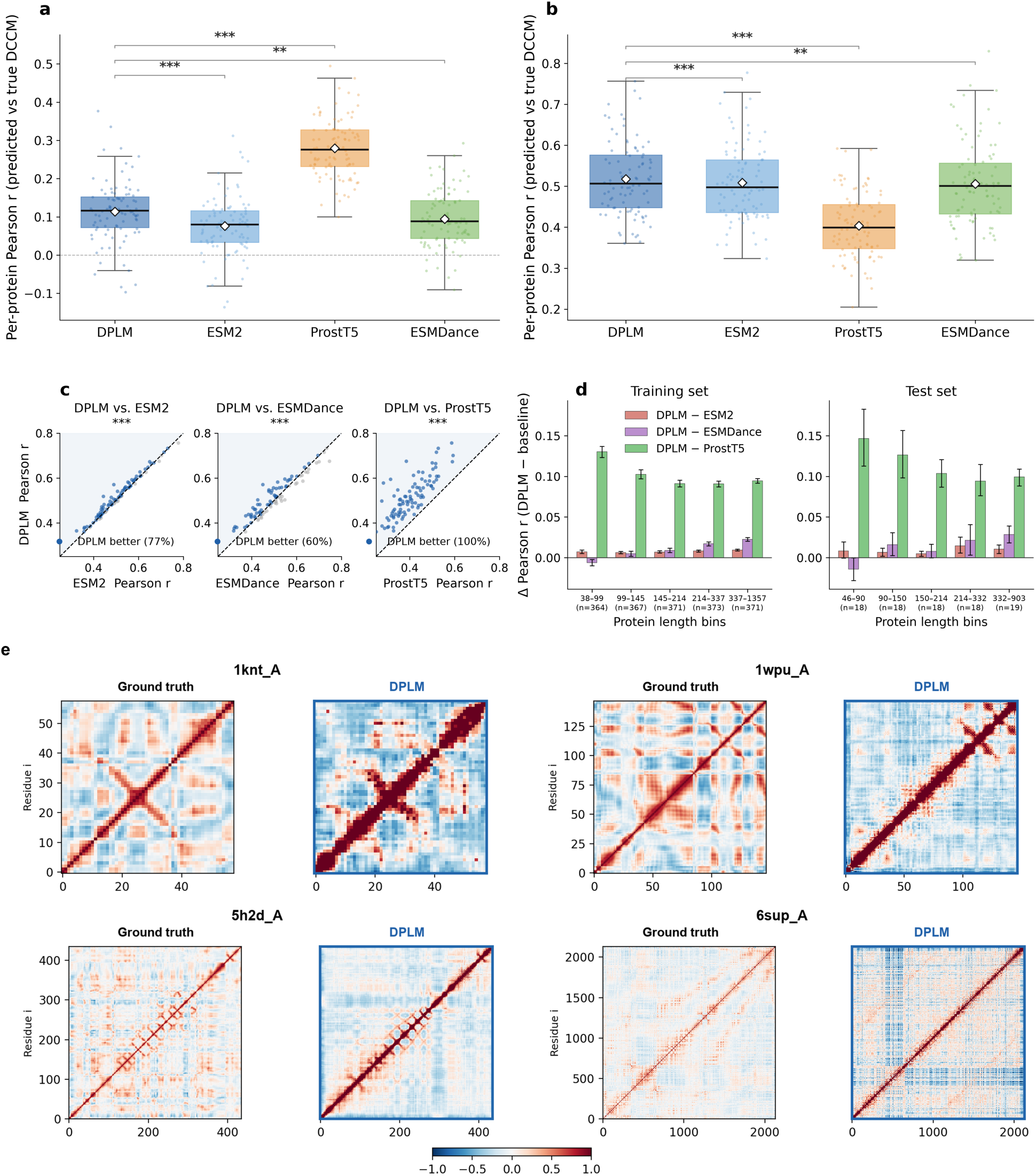
PLM representations recover dynamic cross-correlation between residues. **a.** Zero-shot pairwise cosine similarity of residue embeddings correlated against the DCCM in the Atlas test set. **b.** Zero-shot attention-weighted read-out on the Atlas test set, the same embedding-similarity map gated by each backbone’s own self-attention. **c.** Per-protein scatter of the panel-(b) read-out, DPLM versus each baseline; points above the diagonal (blue) are proteins on which DPLM is better. **d.** The DPLM margin (Δ Pearson r versus each baseline) stratified into protein-length quantile bins, on the training set (left) and the test set (right). **e.** Supervised case studies: a linear regression predictor using the same embedding and attention features, trained on the ATLAS training set and evaluated on held-out proteins. Each block pairs the ground-truth DCCM with DPLM’s supervised prediction. Colour red indicates in-phase motion, blue indicates anticorrelated motion. Predicted maps are per-panel z-scored over the off-diagonal (an affine rescaling that leaves Pearson r unchanged).

Finally, we trained a supervised predictor to align DPLM’s DCCM predictions with ground truth and generalize to unseen proteins. Using the same embedding-and-attention features as in Fig. 3b, we fit a linear regression on the ATLAS training set and evaluated it on held-out proteins (Methods). Figure 3e shows representative predictions across a range of protein sizes, from fewer than 60 to over 2,100 residues. In each case, the predicted DCCM reproduces the salient organization of the ground-truth map. Together, these results show that DPLM’s sequence representation encodes protein dynamics. The pretraining on native MD trajectories helps identify residues that move in a coordinated manner over time.

### DPLM representation leads PLMs on phase-separation clustering and prediction

Liquid–liquid phase separation (LLPS) is a process in which a uniform mixture spontaneously separates into two liquid phases of differing concentration. In cells, LLPS lets proteins and nucleic acids condense into membraneless compartments, such as nucleoli and stress granules^29^. LLPS is intrinsically coupled to protein dynamics, as conformational fluctuations and transient multivalent interactions drive condensate formation and determine their material properties^30^. Dysregulation of LLPS has been implicated in neurodegeneration and cancer^31^, which makes predicting which proteins undergo LLPS of broad biological interest. To test whether our dynamics-aware representation transfers to LLPS prediction, we evaluated DPLM on five curated benchmarks (S1-S5) from a recent comparative study^32^. Each benchmark probes a distinct scenario (see Methods for dataset details).

We first clustered proteins from the embeddings alone. Even without labels, DPLM embeddings organized proteins by phase-separation class. We applied k-means (k = 2) to the t-SNE-reduced embeddings and quantified agreement with the ground-truth partition by the adjusted Rand index (ARI). DPLM gave the strongest clustering on four of the five datasets, while the structure-aware ProstT5 led on the viral set S5 with wide variability (Fig. 4a). This separation emerges without any trained classifier and does not appear in the evolution-based ESM2 baseline, so it can only be explained by the dynamics information that DPLM additionally carries. Supplementary Fig. S6 provides the full unsupervised clustering results and the corresponding embedding visualizations for every model and dataset.

**Fig 4.**
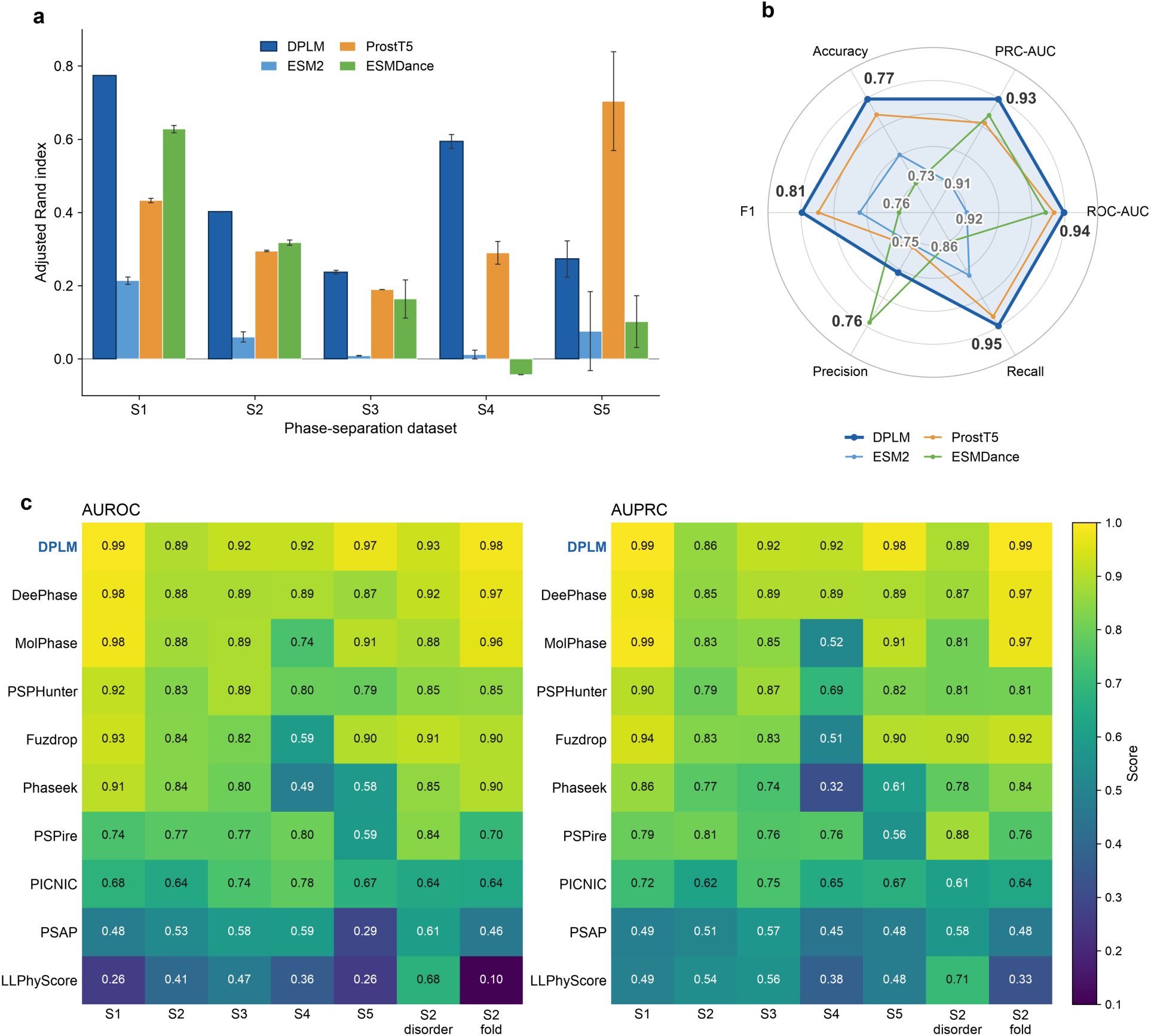
DPLM sequence representations capture liquid-liquid phase-separation propensity. Evaluation on five curated phase-separation benchmarks (S1-S5). DPLM is compared against PLMs (ESM2, ProstT5, ESMDance) and dedicated phase-separation predictors. **a.** Unsupervised agreement between k-means clusters (k = 2, on t-SNE-reduced embeddings) and ground-truth class, measured by the adjusted Rand index. **b.** Supervised classification with an identical classifier head on top of each language model, six metrics averaged over S1-S5 (max and min values for each metric are annotated in the radar plot). **c.** Per-dataset AUROC (left) and AUPRC (right) for DPLM versus nine established phase-separation predictors on S1-S5 and the S2 disordered/folded subsets, rows ordered by mean AUROC.

We then trained a gradient-boosted classifier (XGBoost) on top of each PLM representation (Fig. 4b; classifier details in Methods). DPLM led on all metrics except precision, where it fell marginally behind ESMDance (mean 0.752 versus 0.759). Supplementary Fig. S7 reports the per-method, per-dataset values across S1–S5. To make the evaluation more rigorous and to test whether homologous proteins affect performance, we removed samples in the five datasets with sequence identity > 50% to any sample in the ATLAS training set (Methods), even though DPLM was pretrained in an unsupervised mode. Table S1 lists statistics for the filtered phase-separation dataset, and Fig. S8 shows performance on the filtered datasets. The conclusion does not change on the filtered datasets.

Finally, we benchmarked the same DPLM supervised classifier against the nine published predictors in the review^32^, all of which are dedicated LLPS predictors that do not use PLM embeddings. DPLM ranked first in all cases, leading on both AUROC (0.89 to 0.99) and AUPRC per dataset (Fig. 4c). The largest margins appeared where the original evaluation reported the greatest instability. On folded proteins, where many predictors lose accuracy because they implicitly equate disorder with LLPS propensity, DPLM remained the top predictor, indicating that the dynamics signal is not merely a proxy for disorder. On viral proteins, an under-represented class on which competing methods degrade, DPLM showed no comparable drop (AUROC 0.973). On the LLPS-abolishing-mutation set, where the original study found structure-informed predictors strongest, DPLM still achieved the highest AUROC (0.920, versus 0.802 for PSPire and 0.781 for PICNIC), suggesting that the encoded dynamics carry mutation-sensitive information beyond static structure.

### DPLM embedding complements ProstT5 structure feature in cryptic-site detection

Cryptic binding sites are pockets absent or occluded in the unbound (apo) structure that become druggable only upon a conformational change, which makes them attractive but difficult drug targets^33^. Because a cryptic pocket is defined by motion the apo structure does not display, its detection is intrinsically tied to the conformational ensemble a protein can access. To examine whether DPLM’s dynamics-aware representation carries information useful for this task, we benchmarked DPLM and the baseline PLMs on CryptoBench^34^, a benchmark of cryptic sites annotated from paired apo–holo structures (see Methods for dataset processing). The dataset is imbalanced (positive rate about 5%), as cryptic binding sites are rare. Apart from the input embeddings, which come from different PLMs, all baselines use the same classifier architecture and training protocol (see Methods for training details).

The structure-aware ProstT5 is the strongest overall (Fig. 5a), consistent with the conformational ensemble being the dominant signal for cryptic-site detection. DPLM remains competitive despite using roughly 5× fewer parameters, trailing ProstT5 by only 0.012 in AUC (0.841 vs 0.853), 0.024 in AUPRC (0.321 vs 0.344) and 0.024 in F1 (0.377 vs 0.401) (Fig. 5a). As in the phase-separation task, we ran a sequence-redundancy check and removed proteins with sequence identity > 50% to any protein in the ATLAS training set (Methods). Table S1 presents statistics for the filtered CryptoBench dataset, and the corresponding results in Fig. S9 show that the conclusion still holds.

**Fig 5.**
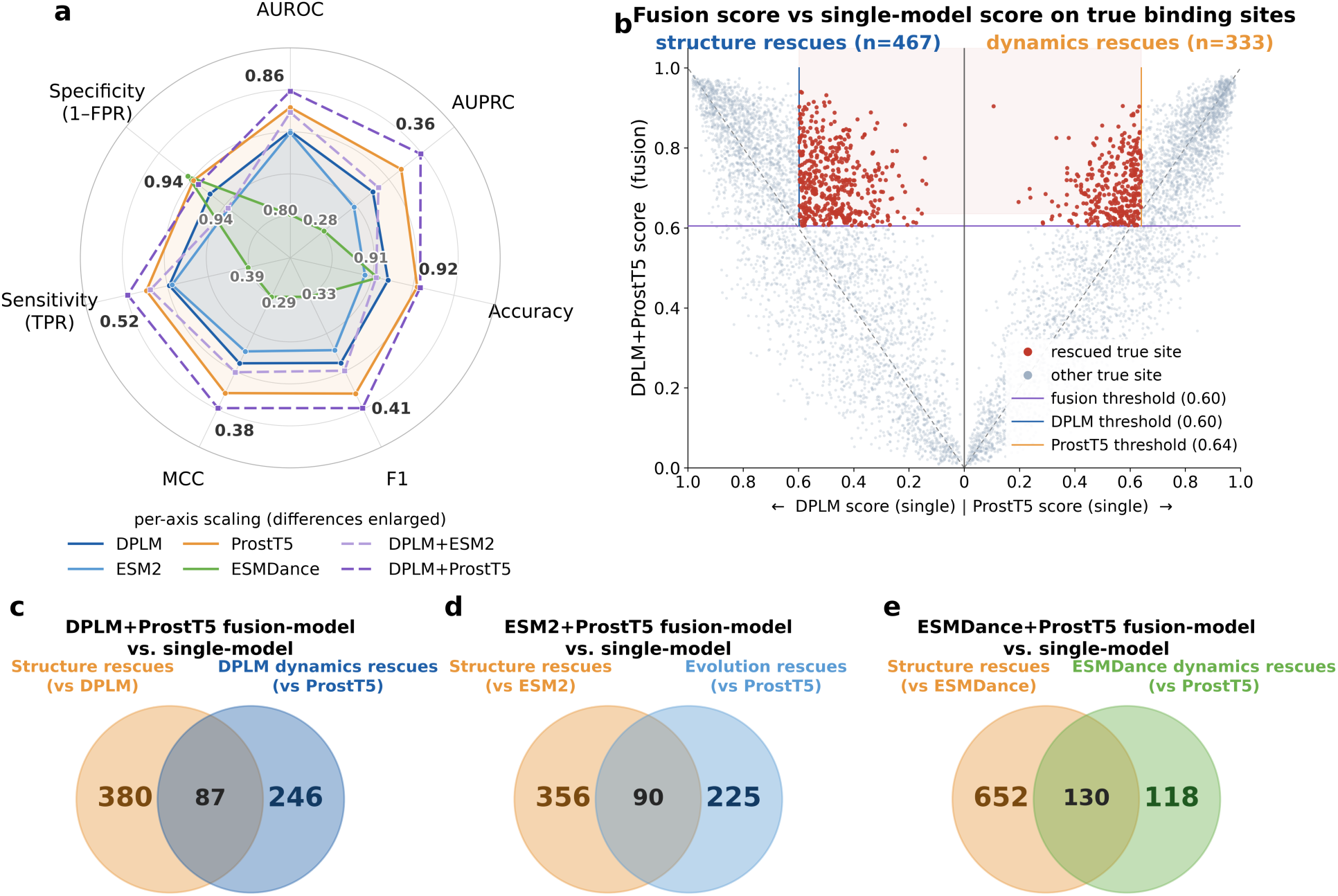
Dynamics and structure contribute complementary residues to cryptic binding-site detection. All models are compared under a fixed classifier and training protocol, varying only the input representation. **a.** CryptoBench cryptic-site detection scored per residue across seven metrics for four single encoders, DPLM, ESM2, ESMDance, and ProstT5, and two feature-fusion classifiers that concatenate DPLM embeddings with ESM2 or with ProstT5 (single models solid, fusions dashed). Each axis is independently scaled to the observed range (per-axis min and max printed at the spoke ends) to make small differences visible. **b.** Back-to-back comparison of the DPLM+ProstT5 fusion against each single model on true binding-site residues: the vertical axis is the fusion score, the left half-axis the DPLM-alone score and the right half-axis the ProstT5-alone score. Red points are true sites the fusion recovers above its operating threshold (horizontal line) that the single model misses. **c.** The overlap between the two rescued-site sets in (b). **d.** Comparing the ESM2+ProstT5 fusion model with the single ESM2 and ProstT5 models, the overlap between the two rescued-site sets. **e.** Comparing the ESMDance+ProstT5 fusion model with the single ESMDance and ProstT5 models, the overlap between the two rescued-site sets.

Because DPLM and ProstT5 encode information from different perspectives (dynamics vs. structure), we tested whether combining them would help. A classifier on concatenated DPLM+ProstT5 features outperformed either single model and every other single representation (AUPRC 0.360, MCC 0.381, F1 0.412; Fig. 5a). The two modalities recover largely distinct residues. The sites rescued by ProstT5 carry a structural signature, whereas those rescued by DPLM do not, suggesting that the dynamics of DPLM rescue them (Fig. S10). Among the true sites rescued by adding one model to the other, only 12% (87 of 713) overlap (Fig. 5b,c). We also trained classifiers on concatenated ESM2+ProstT5 and ESMDance+ProstT5 and derived the overlap between the rescue sets by comparing the corresponding single models (Fig. 5d,e). Adding DPLM rescues more cryptic sites than adding ESM2 or ESMDance, so DPLM is the better complement to a structure-aware model. Supplementary Fig. S11 shows a comprehensive Venn diagram of the overlaps between individual models. On top of ProstT5, adding DPLM introduces the most true positives and the fewest false positives.

### DPLM generalizes to predicting mutation effects on protein thermal stability

Accurate prediction of mutation-induced variance in protein thermal stability is a long-standing and critical problem in protein engineering and design^35^. The effects of single amino-acid substitutions are inherently linked to protein dynamics. A substitution can reshape the protein energy landscape, thereby stabilizing or destabilizing the native state^36^. We therefore applied DPLM to mutation-induced thermal-stability prediction. We first evaluated within-cohort performance on the mega-scale folding-stability dataset^37^. We fit a per-protein linear model, training on 50% of a protein’s mutations and testing on the held-out mutations of the same protein (Fig. 6b,c; dataset and training details in Methods). DPLM, ESM2, and ProstT5 performed similarly and all remained well ahead of ESMDance in both Spearman correlation and mean squared error. Concatenating complementary representations improved within-cohort accuracy further, and using all features (DPLM+ESM-2+ProstT5) reached 0.881, exceeding any single model (Fig. 6 a,b).

**Fig 6.**
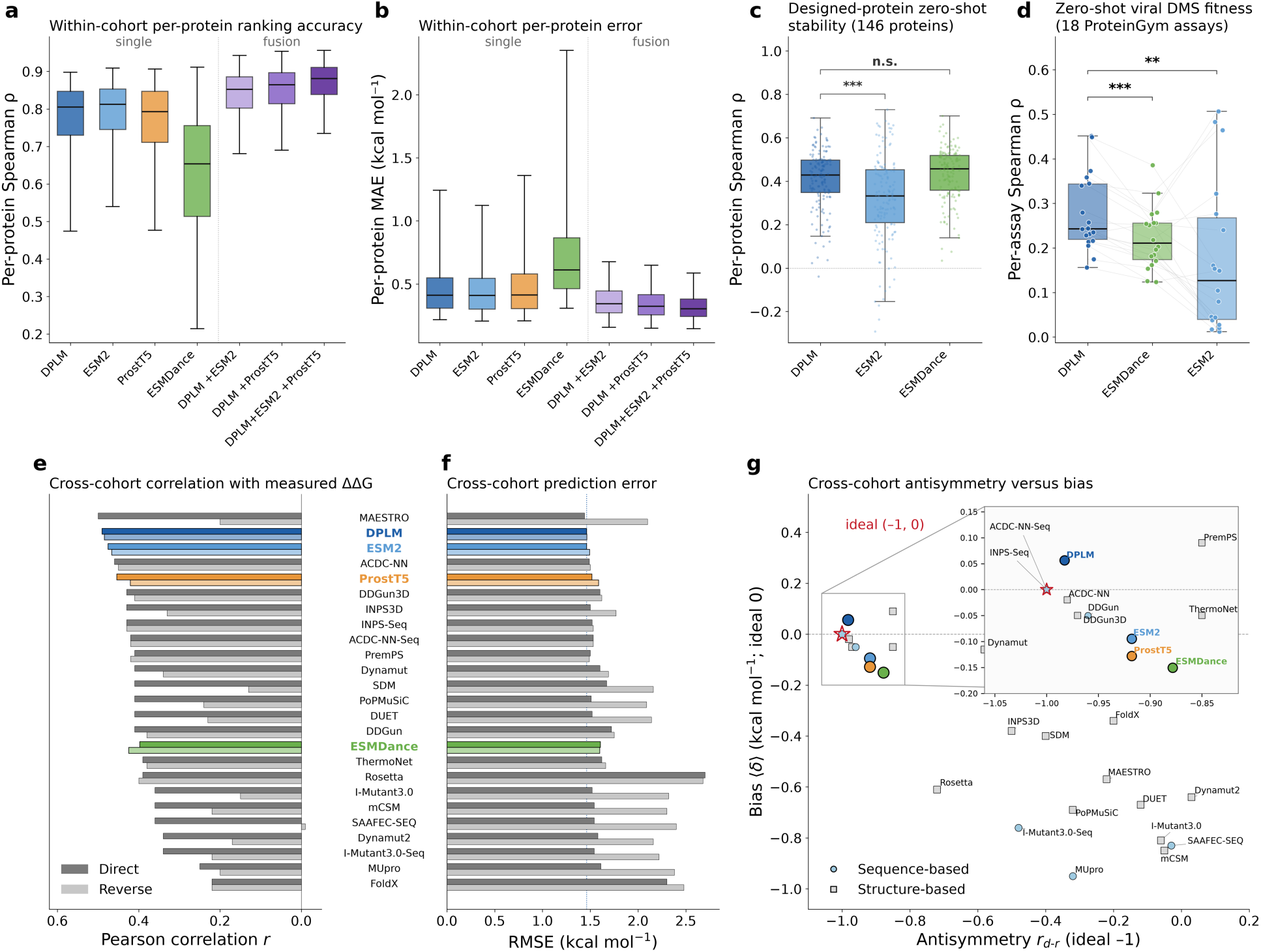
Dynamics-aware sequence representations are robust across mutation-induced fitness change. a, **b.** Within-cohort ΔΔG prediction on the mega-scale folding-stability dataset (412 proteins), scored per protein: **a.** Spearman ρ (ranking; higher is better) and **b.** mean absolute error (kcal mol⁻¹; lower is better), for the four single models and three feature-fusion classifiers (DPLM+ESM-2, DPLM+ProstT5, DPLM+ESM-2+ProstT5). **c.** Zero-shot ΔΔG prediction on the designed proteins (extracted subset from mega-scale dataset). **d.** Zero-shot variant-effect prediction on 18 viral deep-mutational-scanning assays from ProteinGym, scored per assay by Spearman ρ between model score and measured fitness. Each point is one assay, and thin lines connect the same assay across models. **e–g.** Cross-cohort generalization on the independent S669 benchmark, with the four language models placed among published ΔΔG predictors (language models in colour, published tools in grey). **e.** Pearson correlation r and f. RMSE (kcal mol⁻¹) for the direct (darker colour) and reverse (lighter colour) mutation sets; the dotted line in (**f**) marks DPLM’s direct RMSE. **g.** Physical consistency: antisymmetry *r_d_*_-*r*_ (ideal −1) versus mean signed bias ⟨δ⟩ (ideal 0), with the ideal point starred; sequence-based methods are circles, structure-based methods are squares. The inset magnifies the well-calibrated cluster near the ideal point.

A known limitation of ESM2 is that the evolutionary regularities learned from natural protein sequences may not generalize well to out-of-distribution proteins^38^. *De novo* designed proteins may lack close homologs, whereas viral proteins are often highly divergent and underrepresented in general-purpose protein databases^20^. We extracted the designed-protein subset (146 proteins) from the mega-scale dataset and assembled a viral benchmark from ProteinGym^39^ (see Methods for dataset construction details). We then performed zero-shot fitness-effect prediction on both datasets for DPLM, ESM2, and ESMDance. Each variant was scored by its method’s native scheme: ESM2 by the masked-marginal log-likelihood ratio (as in ESM-1v^8^), and ESMDance by its dynamic properties. DPLM used a representation-distance score (wt-mt-RLA^40^) to quantify the variant effect (see Methods for zero-shot scoring function details). On both datasets, the two dynamics-aware methods outperformed ESM2 (Fig. 6c,d). ESMDance led on the designed-protein set by median Spearman correlation, but was not significantly distinguishable from DPLM in distribution. DPLM ranked first on the viral-protein set, significantly ahead of dynamics-supervised ESMDance and ESM2. Dynamics-aware pretraining thus yields sequence representations that generalize better to cases where evolutionary information is weak.

The within-cohort protocol in Fig. 6a,b trains and tests on mutations of the same protein. It primarily measures how well a representation fits proteins already seen during fitting. A more practical and informative test is cross-cohort generalization to proteins absent from the training set. We evaluated this by training on the S8754 dataset^41^ and testing on the independent S669 benchmark^42^ (Fig. 6e-g; dataset and training details in Methods). DPLM reached a direct-mutation Pearson correlation of 0.49, the best among all sequence-based methods and second only to the structure-based MAESTRO (0.50) across the 21 predictors compiled by Pancotti *et al*^42^. Among the four language models evaluated, DPLM led the cross-cohort task (Fig. 6e,f). On reverse mutations, where a model must predict the stabilizing counterpart of each substitution, DPLM retained a correlation of 0.48, the highest of any method in the comparison, whereas most predictors dropped sharply (for example, MAESTRO from 0.50 to 0.20). As with the other downstream tasks, we excluded proteins in the mega-scale, ProteinGym viral, and S669 sets with sequence identity > 50% to any protein in the ATLAS training set. Table S1 gives the filtered dataset statistics, and Figs. S12–S14 show the filtered results. All conclusions from the full datasets still hold for the filtered datasets. No protein is filtered from the designed-protein dataset, as all proteins are already below the identity threshold.

We further quantified physical consistency with a previously defined antisymmetry index: *r_d_*_-*r*_ (Equation (6) in Methods). Additionally, to measure the average bias toward a specific class, we adopted a bias score 〈*δ*〉 (Equation (7) in Methods). The DPLM-based method has near-ideal antisymmetry (−0.98; ideal −1) and negligible bias (0.05 kcal mol⁻¹; ideal 0). This places the DPLM-based method among the best-calibrated predictors of any class, despite using only sequence input (Fig. 6g). Across the four PLMs, DPLM sits closest to the ideal (−1, 0) corner of this antisymmetry–bias space (Fig. 6g). ESM2, ProstT5 and ESMDance all show a larger destabilization bias (−0.09, −0.13 and −0.15 kcal mol⁻¹, respectively) and weaker antisymmetry (−0.92, −0.92 and −0.88, respectively). Among the task-specific methods, ACDC-NN-Seq and INPS-Seq hard-code antisymmetry directly into the model architecture.

## Discussion

DPLM is a general-purpose representation model. Its dynamics-aware embeddings transfer across downstream tasks and improve prediction of dynamics-related properties. Unlike previous work, the dynamics in DPLM does not come from a conformational ensemble or from predefined dynamic descriptors. A conformational ensemble captures the diversity of conformational states but does not necessarily encode their temporal ordering or transition kinetics^43,44^. Predefined descriptors fix in advance which aspects of the dynamics are kept and reduce each trajectory to a set of summary statistics. Instead, DPLM embeds time-resolved molecular motion directly from MD trajectories: it treats each trajectory as a video of Cα contact maps and encodes it with a pretrained ViViT video model, so the joint spatiotemporal structure of the motion is preserved. The sequence and MD embeddings are aligned in a shared latent space and are not tied to any specific feature. The resulting representation can still be quantified against specific features, showing high correlation with RMSF (Fig. 2) and DCCM (Fig. 3). Learning the geometry of motion, rather than fitting a chosen summary of it, therefore appears to retain more transferable dynamic information.

The advantages of DPLM on phase-separation and thermal-stability prediction can be understood from the physical principles underlying these properties. LLPS is driven by networks of weak, multivalent intermolecular interactions, frequently involving intrinsically disordered regions^29^. The inherent dynamics of these regions directly influence the interaction patterns that promote or oppose phase separation^29,45,46^. Thermal stability is likewise intimately coupled to molecular motion. A folded protein is stabilized by a cooperative network of residue–residue interactions, and mutations that alter packing or interaction networks can consequently perturb protein dynamics and shift the free-energy balance between folded and unfolded states^47,48^. By learning residue-level fluctuations and coordinated motions from physical MD trajectories, DPLM captures dynamics-features related to these processes rather than inferring them solely from sequence or a single static structure. This may explain its gains in LLPS and thermal stability prediction, particularly for proteins with weak evolutionary priors, including those absent from pretraining, *de novo* designed proteins, and viral sequences with sparse homology (Figs. 4 and 6).

Cryptic binding-site prediction provides a complementary case in which both static geometry and dynamics are important. Cryptic pockets are absent or occluded in experimentally observed apo structures and become accessible through conformational rearrangements^33,49^. Their formation is therefore inherently dynamic, and atomistic simulations have been widely used to characterize such pocket-opening events^33,50^. However, these transitions can be rare and separated from the native basin by substantial free-energy barriers. DPLM is trained on relatively short equilibrium trajectories from ATLAS, which primarily capture fluctuations around native conformations and may therefore undersample activated pocket-opening events. This provides a physical explanation for its weaker performance on this highly localized task. The 3B-parameter structure-trained ProstT5 achieves the strongest individual performance (Fig. 5a), although the 650M DPLM remains close, trailing by only 0.012 in AUC and 0.024 in F1. Importantly, their errors are complementary: combining DPLM with ProstT5 recovers cryptic-site residues missed by either representation alone and outperforms every individual method (Fig. 5). These results suggest that static structural information and molecular dynamics capture complementary aspects of cryptic-pocket formation: structure constrains the local geometry from which a pocket can emerge, whereas dynamics describes the motions through which that geometry becomes accessible.

The pattern across tasks points to a broader methodological observation. Unsupervised and zero-shot evaluations may reveal differences in representation quality more directly than supervised benchmarks^51,52^. When we read out the pretrained representations without task-specific training, DPLM consistently separates from the baselines. This pattern appears in RMSF tracking, zero-shot DCCM recovery, unsupervised clustering of phase-separation classes, and zero-shot viral protein fitness prediction. In supervised settings, a trainable prediction head introduces additional capacity that can compensate for differences in the underlying representations. As a result, the performance gap between methods often narrows. These findings highlight the value of unsupervised and zero-shot evaluations for assessing the information encoded directly in pretrained protein representations^53^.

Two limitations define the scope of this study and suggest directions for future work. First, DPLM is trained on a modest amount of MD data from the ATLAS database. This limits the diversity of molecular motions available during training. Larger datasets and longer simulations could expose the model to a broader range of dynamical processes. Data consistency may also be important. Adding mdCATH trajectories to the training set did not improve performance (Fig. S1). One possible explanation is the difference in simulation protocols and force fields between ATLAS and mdCATH, which may introduce heterogeneity into the dynamics used for training^54^. Second, the current MD encoder limits model scalability. The ViViT encoder is the base google/vivit-b-16×2-kinetics400 model (≈86M parameters), the only pretrained ViViT checkpoint readily available. Its fixed input resolution limits how much of a trajectory can be represented at once. Future work should therefore expand high-quality MD training data and develop scalable trajectory encoders that capture longer and more diverse molecular motions. Evaluating DPLM on a broader range of datasets and downstream tasks is a further priority, to establish how widely the dynamics-aware representation generalizes.

## Methods

### DPLM architecture

#### Sequence encoder

The sequence branch is built on ESM2 (650M parameters; esm2_t33_650M_UR50D), a 33-layer transformer with embedding dimension 1,280^6^. The pretrained weights are kept frozen. To adapt the model to the dynamics-alignment objective while preserving its pretrained knowledge, we insert Houlsby-style bottleneck adapters^24^ into the top 10 transformer layers (layers 24-33), placing one adapter after the multi-head-attention sublayer and one after the feed-forward sublayer in each of these layers. Each adapter is a residual MLP that projects the 1,280-dimensional hidden state down through a bottleneck, applies a ReLU nonlinearity, projects back up, and adds the result to the input, followed by layer normalization. Only the adapter parameters are trainable in this branch. Sequences are truncated to a maximum length of 512 residues. A protein-level sequence embedding is obtained by masked mean-pooling of the final-layer residue representations over non-padding positions.

#### MD encoder

Each MD trajectory is encoded as a sequence of residue-residue contact maps. For each frame, we compute the Cα-Cα distance matrix using MDTraj^55^, normalize it to its maximum value, and replicate it across three channels to form an image. Frames are tiled to 224×224 with a fragment length of 224 residues and a 30-residue overlap for longer proteins, yielding a contact-motion video. Videos are grouped into non-overlapping 32-frame clips and encoded with a Video Vision Transformer (ViViT, google/vivit-b-16×2-kinetics400)^23^, taking the [CLS] token (768-dimensional) of each clip. Clip embeddings are mean-pooled into a single 768-dimensional trajectory representation per protein. The ViViT backbone is frozen; only its downstream projection is trained. Trajectory features are precomputed once and cached, so ViViT is not evaluated during contrastive training.

#### Projection heads

Each branch is followed by a projection head implemented as a batch-normalized MLP. The protein-level head is a two-layer MLP with an inner dimension of 4,096, projecting to a 1,024-dimensional space; a separate single-layer residue-level head projects to 64 dimensions. Contrastive alignment is performed in the 1,024-dimensional protein-level space.

### Contrastive pretraining

The projected embeddings from sequence and MD branches are aligned through a multi-view contrastive learning objective. The training objective encourages embeddings from the same protein (sequence embedding *E_i_^seq^* and MD embedding *E_i_^md^*) to be closely aligned, while simultaneously pushing apart embeddings from different proteins (de-alignment). This multi-view setting ensures that both modalities capture protein-specific information while avoiding collapse to trivial representations. The contrastive objective is based on a modification of the NT-Xent loss introduced in SimCLR^25^. We modified it to address cross-view contrast as introduced in X-CLIP^27^. For a positive pair (*E_i_^seq^*, *E_i_^md^*) from protein i, the loss is defined as:

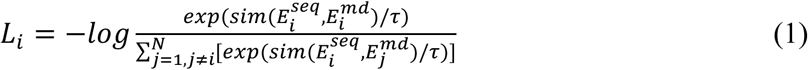

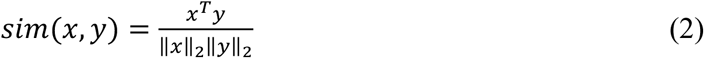

where *τ* is a temperature parameter (set as 0.1), N is the batch size. This formulation maximizes similarity between paired sequence/MD embeddings from the same protein while minimizing similarity between embeddings from different proteins. The total contrastive loss is computed across all positive pairs in each minibatch.

### DPLM Training data and optimization

We train on all-atom MD trajectories from the ATLAS database^26^, using three independent simulation replicas per protein. There are 1938 proteins in the current version of ATLAS database. We select about 5% samples (92) as the test set with the following constraints: 1) The test proteins have different ECOD domains^56^ from the training set proteins, so the two are structurally non-redundant, 2) The test proteins have a spread protein sequence length, and 3) The test proteins have protein sequence identity less than 30% with the training set proteins. The model is optimized with Adam (learning rate 5×10⁻⁵ for both branches, β₁ = 0.9, β₂ = 0.999, ε = 10⁻⁸, weight decay 5×10⁻⁴) for 30,000 steps with a batch size of 32, a 500-step linear warm-up followed by cosine decay of the learning rate to a floor of 10⁻⁶, gradient clipping at norm 1.0, and bf16 mixed precision. A fixed random seed is used for reproducibility.

### Residue flexibility and embedding analysis

For the residue-level flexibility analysis (Fig. 2a-b, e) we reduced each residue’s high-dimensional embedding to a single scalar. Given the per-residue representation ***e****_i_* ∈ ℝ*^D^* for residue *i* (D = 1,280 for the ESM2-based models), we defined the residue embedding scalar as its Euclidean (L2) norm:

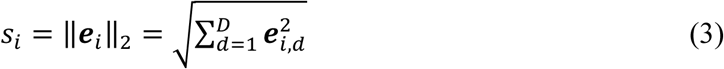

a single positive value summarizing the magnitude of the residue’s embedding. We then measured the association between this scalar and per-residue mobility, quantified by the molecular-dynamics root-mean-square fluctuation (RMSF) using Spearman’s rank correlation. The RMSF values were computed from the ATLAS test trajectories, defined as:

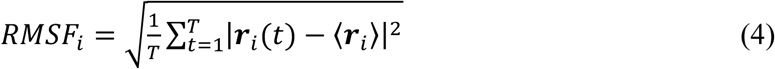

Where ***r****_i_*(*t*) is position of residue *i* at frame 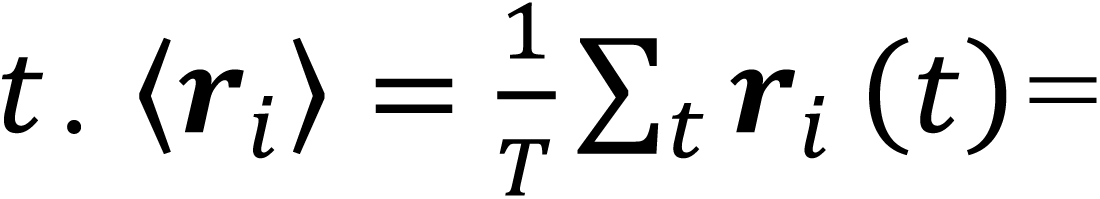 time-averaged position of residue *i*. *T*= total number of trajectory frames.

Both the residue embedding scalar and the RMSF values were pooled over all residues of all test proteins (whole-dataset correlation, Fig. 2a) and computed within each protein and summarized as a distribution across proteins (per-protein correlation, Fig. 2c). For two of the baselines (ProstT5 and ESMDance) the raw scalar anti-correlates with flexibility; because the quantity of interest is the strength of the scalar-flexibility association rather than its sign, we report the magnitude of the correlation (equivalently, sign-flipping the scalar to positive) so that all methods are compared on the same footing. The main-text figure is based on a single replicate, and the corresponding results for all three replicates are provided in Supplementary Figs. S1 and S2.

Ground-truth DCCMs were computed from the all-atom ATLAS MD trajectories. For a pair of residues *i* and *j* it measures the normalized correlation of their positional fluctuations over a molecular-dynamics trajectory, defined as:

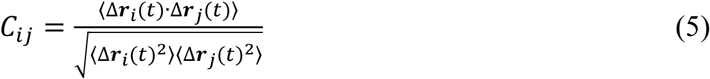

Where Δ*r_i_*(*t*) = ***r****_i_*(*t*) − 〈***r****_i_*〉, the instantaneous displacement of atom *i*from its mean position. The averages ⟨⋅⟩ are taken over time/frames. *C_i_*_j_ ∈ [−1,1], +1 = fully correlated motion, −1 = fully anti-correlated, 0 = uncorrelated. ESMDance was trained to predict RMSF and DCCM directly. We exclude that supervised read-out here to avoid information leakage, since ESMDance uses the ATLAS dataset for training. Our focus is whether dynamics is present in the representation itself, because that representation is what downstream tasks consume. We therefore extracted, processed and evaluated the embeddings from all methods in the same way.

Model performance was quantified as the per-protein Pearson (and Spearman) correlation between the upper triangles of the predicted and ground-truth matrices. We evaluated three read-outs of increasing supervision, all built on frozen backbones (DPLM, ESM2, ProstT5 and ESMDance). (i) Zero-shot cosine read-out. The pairwise cosine-similarity matrix of per-residue embeddings was correlated with the DCCM with no fitted parameters. (ii) Zero-shot attention-weighted read-out. Each backbone’s self-attention was read through its own API, symmetrized, head-averaged and averaged over all layers into a single pairwise map *A*; the read-out *A*⨀*cos*(*e_i_*, *e*_j_), attention gating the embedding-similarity signal, was correlated against the DCCM, again with no fitted parameters. (iii) Supervised predictor. The predictor combines two pathways. A per-residue pathway projects each residue embedding through a two-layer MLP (hidden dimension 512, projection dimension 128, dropout 0.2) and takes the cosine similarity between projected residues, giving a score *S_emb_*[*i*, *j*] for every residue pair. An attention pathway reads the backbone’s own self-attention. For each pair, a learned linear read-out maps the C head-averaged, symmetrized attention channels to a single value *a*[*i*, *j*]. The two pathways fuse multiplicatively as *tan*ℎ(*S_emb_*⨀(1 + *a*)), so attention gates the per-residue score. The attention read-out is zero-initialized, and training therefore starts from the pure per-residue predictor and learns the attention contribution from there. The comparison holds the attention-channel count equal across backbones. The backbones differ in depth, 33 layers for DPLM and ESM2, 24 for ProstT5, 12 for ESMDance, so we take the last ten attention layers of each, ten being the largest count the shallowest backbone (ESMDance) supports. Every method therefore contributes the same C = 10 channels. Both the embeddings and the attention channels come from a single frozen forward pass per protein. We trained the predictor on the ATLAS training set with Adam (learning rate 1×10⁻³, weight decay 1×10⁻⁵, 40 epochs, 10% validation split). The loss combined the mean-squared error and a correlation term on the DCCM upper triangle, and we kept the checkpoint with the best validation correlation. We repeated training over three seeds and took each per-protein score as the mean over seeds. Evaluation used the held-out ATLAS test proteins.

### Phase separation prediction

The five evaluation datasets (S1–S5) are taken from the comparative evaluation of Gilroy & Barr^32^, each probing a distinct scenario in which LLPS predictors are known to succeed or fail. Throughout, positives are experimentally validated phase-separating proteins and negatives are sequences considered unlikely to phase separate; the sets differ in the structural and biological character of their proteins. Standard (S1): a balanced, general-purpose set of 200 proteins (100/100) representing the conventional case, validated LLPS drivers versus well-folded, low-disorder proteins with resolved structures. Disordered vs. folded (S2): the same distinction on 400 proteins (200/200), each additionally labelled disordered or folded, yielding two 200-protein subsets evaluated separately (Fig. 3b) to test whether a predictor identifies phase-separating folded proteins and non-phase-separating disordered ones rather than defaulting to “disordered = LLPS.” High-confidence benchmark (S3): a stringently curated set of 200 proteins (100/100) minimising false negatives, the central difficulty in LLPS prediction, by restricting positives to high-confidence driver-exclusive phase-separating proteins. Mutations (S4): 36 sequences, twelve phase-separating wild-types paired with 24 point-mutant variants shown to abolish phase separation, testing sensitivity to the small sequence changes that switch phase behaviour off. Virus (S5): 60 viral proteins (30/30), a class under-represented in training data, measuring generalization beyond the human-centric proteins most models are trained on.

Each protein sequence was encoded with the pretrained DPLM sequence encoder, and the residue-level representations (excluding the terminal tokens) were mean-pooled into a single 1,280-dimensional protein embedding. A gradient-boosted decision-tree classifier (XGBoost) was trained on these embeddings to distinguish phase-separating from non-phase-separating proteins, using the MolPhase training set^57^ (606 positive and 1,362 negative sequences). The classifier used 500 trees with maximum depth 6, learning rate 0.05, subsampling and column-subsampling of 0.8, and a class-imbalance weight set to the negative-to-positive ratio; it was optimized with the logistic-loss objective. The trained model was then applied without further tuning to the five evaluation datasets (S1-S5), and performance was quantified by AUROC and AUPRC against the nine predictors compiled by Gilroy & Barr^32^. To visualize the embedding space, protein embeddings were standardized, reduced to 50 principal components, and projected to two dimensions with t-SNE (perplexity 30); unsupervised k-means clustering (k = 2) was applied to the standardized embeddings, and agreement with the experimental labels was measured by the adjusted Rand index.

### Cryptic binding-site prediction

We evaluated residue-level cryptic-site prediction on CryptoBench^34^, a dataset of cryptic binding sites annotated from paired apo-holo structures. Each apo protein is labelled at the residue level: residues belonging to any experimentally observed cryptic pocket (unioned across all holo partners of that apo structure) are positives and all remaining residues are negatives, yielding a strongly imbalanced per-residue binary classification task. We used the benchmark’s predefined training folds and held-out test split (test set with 222 apo structures and a train set with 885 apo structures), so that no test protein contributes to model fitting or threshold selection.

For every apo chain, we extracted the observed-residue sequence from its structure and computed per-residue embeddings with each protein encoder under comparison. For DPLM this yields an L×1280 matrix per chain from the ESM2 650M backbone with Houlsby adapters (L×2560 for the ESM2 3B variant); ESM2, ESMDance and ProstT5 embeddings were produced analogously at their native dimensionalities. Because CryptoBench pocket annotations use author residue numbering, which does not coincide with the embedding index whenever residues are unresolved in the structure, we stored an explicit residue-identifier sidecar for each chain and mapped every annotated pocket residue to its embedding row through that index; multi-chain apo entries were handled by concatenating per-chain sequences under composite chain-residue keys. Sequences were truncated at 1,022 residues (the ESM2 context limit).

To isolate the contribution of the representation, we held a single classifier and training protocol fixed and varied only the input embeddings. The classifier is a gradient-boosted decision-tree model (XGBoost); because a tree ensemble is position-independent, each protein’s L×D block of per-residue embeddings was flattened into a single [N_residues, D] feature matrix and every residue was classified from its own embedding vector, with residue order carrying no information.

The strong class imbalance (∼5% cryptic residues) was addressed by weighting the positive class with scale_pos_weight = n_negative / n_positive. The model used 500 trees, maximum tree depth 6, learning rate 0.05, subsample and column-subsample fractions of 0.8, minimum child weight 1, L2 regularization 1.0 and a logistic (log-loss) objective (random seed 42). Following the benchmark protocol, we used the predefined training folds in 4-fold cross-validation to select the decision threshold that maximizes F1, then trained a final model on the full training set and evaluated it once on the held-out test set. We report area under the ROC curve (AUC), area under the precision-recall curve (AUPRC), accuracy, true-and false-positive rates (TPR, FPR), Matthews correlation coefficient (MCC) and F1; given the strong class imbalance, AUPRC, MCC and F1 are the primary metrics.

### Zero-shot viral fitness prediction

We evaluated zero-shot variant-effect prediction on viral deep-mutational-scanning assays from ProteinGym^39^. From the 217 substitution assays, we kept the 31 annotated as viral, then removed six Tsuboyama_2023 folding-stability assays on small phage and viral domains and two lambda-repressor assays that report expression rather than fitness, and dropped five assays with wild-type length above 1,024 residues (the ESMDance threshold, above which its pairwise dynamic properties no longer fit in memory), leaving 18 assays and 178,503 single mutations. Each method was scored with the read-out its model family is designed for, so the comparison is not confounded by an ill-suited metric. For DPLM, we used a representation-distance score (wt-mt-RLA^40^): each variant is scored as the negative log of the mean per-residue Euclidean distance between the mutant and wild-type final-layer residue embeddings (terminal tokens excluded), so that smaller representational perturbation implies higher predicted fitness. For ESM2, we used masked-marginal scoring: at each mutated position, the residue is masked, and the variant score is the sum over mutated sites of the log-likelihood ratio *logP(mutant) − logP(wild type)* under the masked-language-model head. For ESMDance, we used its native dynamic-property read-out: for each of its predicted dynamic properties we took the mean absolute relative change between wild-type and mutant, quantile-normalised these across an assay’s mutants and combined them by geometric mean, negating the result so that larger predicted dynamic disruption corresponds to lower fitness. Variant scores were ranked against measured DMS fitness by Spearman correlation per assay, and methods were compared by paired Wilcoxon signed-rank tests across the 18 assays.

### Within-cohort ΔΔG prediction (mega-scale)

We applied the DPLM embeddings to the mega-scale protein folding-stability dataset^37^, which includes ΔΔG values for 379,865 single mutations and 147,926 double mutations across 412 proteins, encompassing both natural and designed proteins. Each wild-type and mutant sequence was encoded with the pretrained DPLM sequence encoder and mean-pooled into a 1,280-dimensional embedding, and the feature for each mutant was the difference between its embedding and that of the corresponding wild type. To align these features with ΔΔG, we fitted supervised linear-regression models. For each protein, 50% of mutations were randomly sampled for training and the remaining 50% were used for testing; performance was summarized by the per-protein Spearman correlation and mean absolute error. The identical protocol was applied to ESM2, ESMDance and ESMDance embeddings for comparison.

### Zero-shot *de novo* designed protein ΔΔG prediction (mega-scale)

We extracted the de novo designed protein set from the mega-scale dataset. The designed protein set comprises 123,099 single-and multi-point substitutions over 146 wild types. The zero-shot prediction protocol is the same as for the viral proteins.

### Cross-cohort ΔΔG prediction

To test generalization to unseen proteins, we trained a mutation-site-aware regressor on the frozen DPLM sequence encoder (ESM-2 650M, inputs capped at 512 residues), augmented with Houlsby bottleneck adapters in its top layers: the 10 adapters carried over from DPLM pretraining were held frozen, and 4 new trainable adapters were introduced on top, so that only these 4 adapter layers and the MLP head were learned. For a wild-type/mutant pair, the encoder representations of both sequences are combined into a feature vector concatenating the difference at the mutated residue positions, the difference in masked whole-protein mean pooling, and the two site representations; an MLP head (LayerNorm, hidden dimension 512, GELU, dropout 0.2) then outputs ΔΔG. The model is direction-aware rather than antisymmetric by construction: during training, each mutation is augmented with its reverse (mutant to wild-type) counterpart under a sign-flipped label, so the physical relation ΔΔG(wt to mut) = −ΔΔG(mut to wt) is learned from data (test mutations are never augmented). To assess the antisymmetric property of predictors, we adopted the previously defined index *r_d_*_-*r*_, defined as:

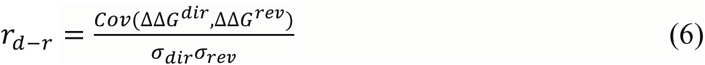

Where Cov is the covariance and *σ* is the standard deviation. To measure the average bias toward a specific class, we adopted the bias score 〈*δ*〉, defiend as:

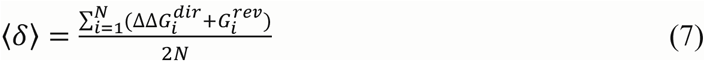

A perfectly antisymmetric and unbiased method should have *r_d_*_-*r*_ equal to −1, whereas 〈*δ*〉 equals to 0.

We trained on S8754 dataset from Xu et al. work^41^. The dataset contains 8,754 single-point mutations across 301 proteins, aggregated from ProThermDB^58^ and ThermoMutDB^59^. S8754 fully contains the S2648 set^60^, with entries >25% sequence identity to the test set removed, and evaluated on the independent S669 benchmark^42^ (669 single-point mutations, 94 proteins). Training used a Huber loss (β = 0.5) and the Adam optimizer with decoupled weight decay (learning rate 3×10⁻⁵, weight decay 5×10⁻³, ε = 1×10⁻¹⁶, gradient clipping 1.0) for 30 epochs at batch size 32 in bf16, with a 300-step warmup and cosine decay to 1×10⁻⁶; the protein-wise train/validation split follows the existing pipeline (test_size 0.2, seed 42). On S669, we report the Spearman and Pearson correlation, RMSE and MAE between predicted and measured ΔΔG for both direct and reverse mutations, together with the antisymmetry correlation *r_d_*_-*r*_ and mean bias 〈*δ*〉; baseline values are taken from an evaluation work by Pancotti *et al.*^42^, all methods trained on thermodynamic ΔΔG data and evaluated on the same S669 test set.

### Sequence-redundancy filtering of evaluation sets

To prevent DPLM from being scored on proteins it had memorized, every evaluation set was screened for sequence redundancy against the ATLAS training split and redundant samples were removed. The reference database was the 1,846 training proteins of the ATLAS. For each benchmark we extracted the sequence of every evaluation unit (the DMS target sequence for ProteinGym viral, the wild-type sequence for the ddG sets, the apo chain sequence for CryptoBench, and the individual sequences of the phase-separation test tables), and searched them against the ATLAS training database with MMseqs2 v18.8cc5c (easy-search, --min-seq-id 0.30-c 0.5 --cov-mode 1-s 7.5 --max-seqs 4000, output query,target,fident,alnlen,qlen,tlen). The prefilter was deliberately set at 30% identity, below the 50% removal threshold, so that borderline alignments were still measured rather than discarded before scoring. For each alignment we computed sequence identity over the shorter of the two sequences (the CD-HIT convention), pid_short = 100 × fident × alnlen / min(qlen, tlen), and assigned each evaluation sequence the maximum pid_short over all of its hits. Any evaluation sequence whose best pid_short exceeded 50% was removed, so that every retained sample is below 50% identity to all 1,846 ATLAS training proteins. Filtering was applied only to evaluation data, the training partitions of the downstream benchmarks (S8754 for ddG S669, the CryptoBench train folds, the Molphase phase-separation training tables) were left untouched, since redundancy there does not inflate the reported scores. Each filtered set was written back in the exact file format its benchmark consumes, so no evaluation code was modified.

### Data, Materials, and software availability

The data used and processed for evaluation and training are available via Hugging Face at https://huggingface.co/datasets/Yuexuhug/DPLM/tree/main. Except for three large-scale datasets, which can be accessed through their original site: Atlas MD trajectories and dynamics property analysis at https://www.dsimb.inserm.fr/ATLAS/index.html; CryptoBench dataset + mmCIF structures at https://osf.io/pz4a9/files/osfstorage; Mega-scale ΔΔG Tsuboyama2023_Dataset2_Dataset3_20230416.csv at https://zenodo.org/records/7992926. The GitHub repository for the project is at https://github.com/yuexujiang/DPLM_release. The trained model checkpoints are available via Hugging Face at https://huggingface.co/Yuexuhug/DPLM/tree/main.

## Acknowledgements

The authors gratefully acknowledge National Institutes of Health (R01LM014510, R35-GM126985) and National Science Foundation (2434170) for funding this project.

