## supplementary information for "DPLM: Dynamics-aware Protein Language Model via Contrastive Learning between Sequence and Molecular Dynamics Simulation Trajectory"

### This PDF file includes:

Supplementary Figures S1-S14

Supplementary Table S1

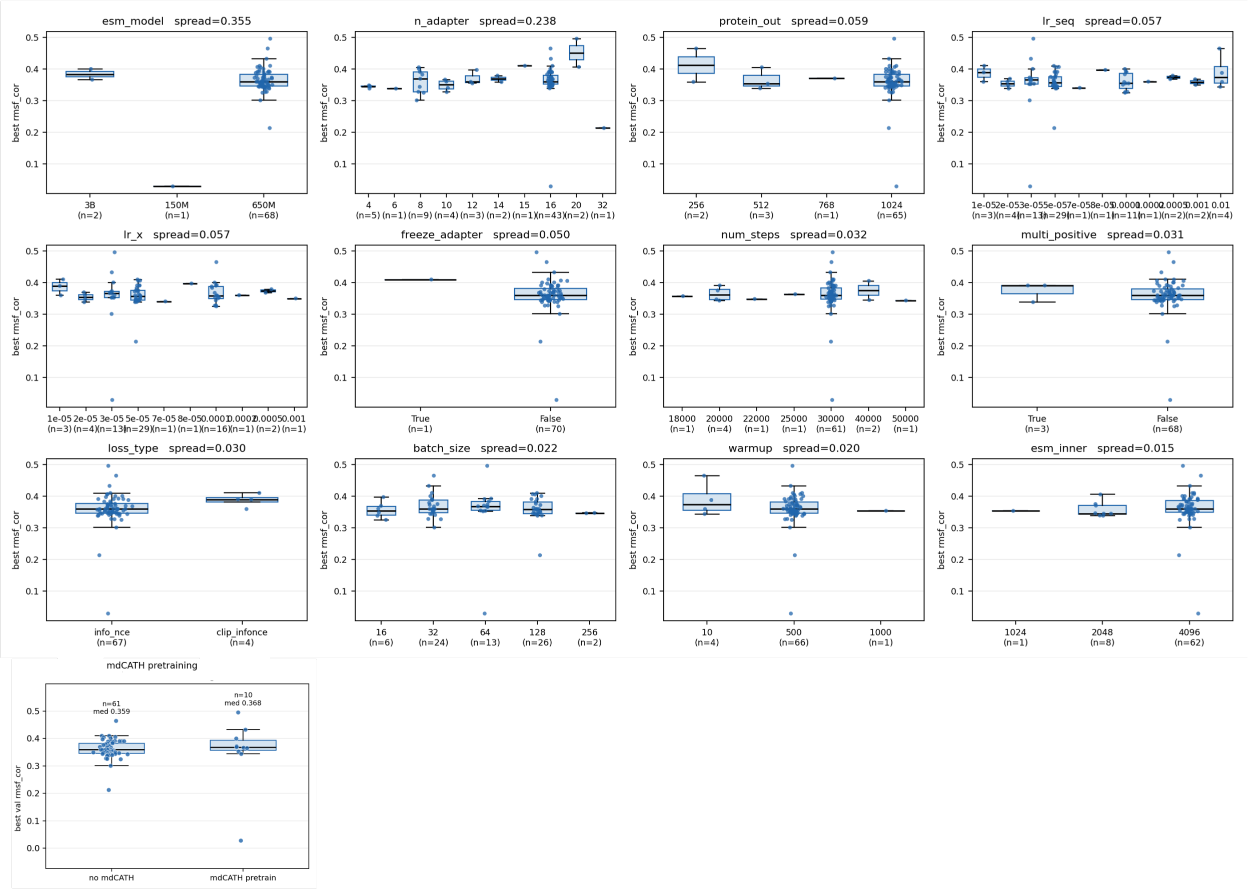


**Fig S1. Marginal effect of each training hyperparameter on validation RMSF correlation.** Each panel shows one hyperparameter. The results are confounded, runs differ in many hyperparameters at once. Ranked by spread of group medians.

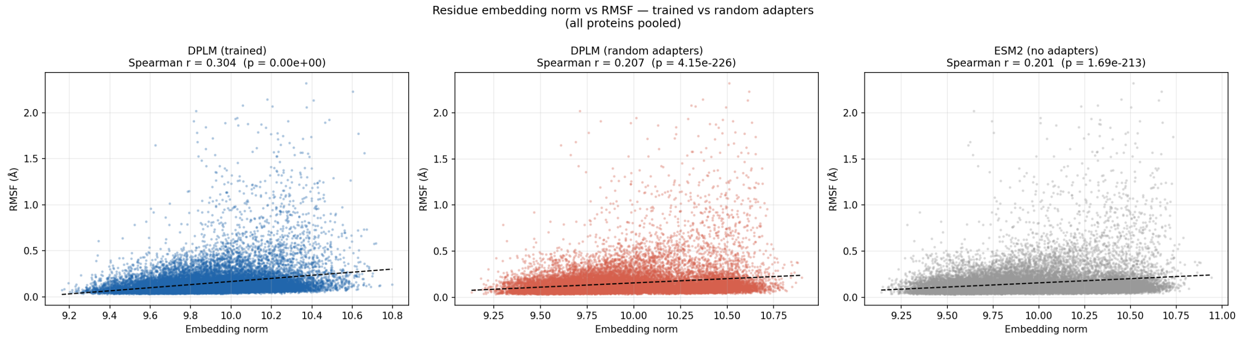


**Fig S2. The residue embedding scalar vs RMSF** from **a.** Trained DPLM, **b.** DPLM without training (randomly initialed adapter weights), and c. native ESM2 650M model. The result is based on replicate 1. It indicates that the adapters in DPLM play a essential role in dynamics learning.

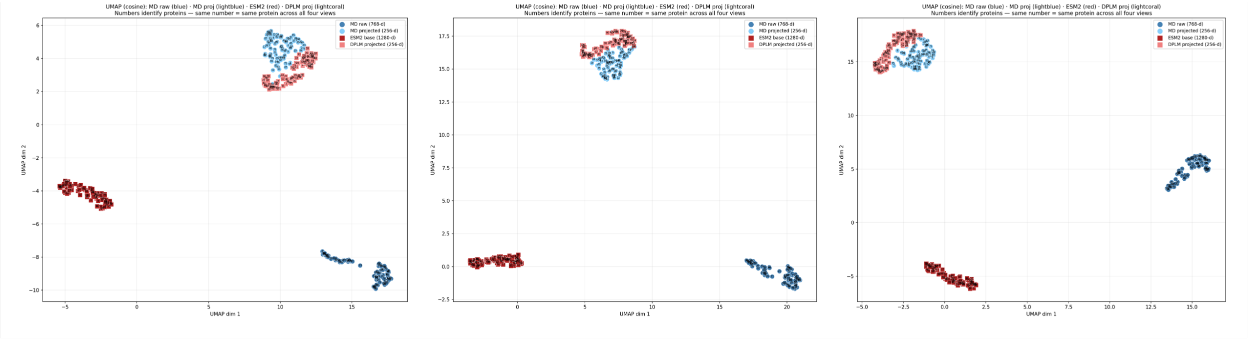


**Fig S3. Sequence–MD embedding alignment before and after contrastive learning.** Two-dimensional joint cosine-UMAP of four embedding views for each ATLAS test protein, shown for the three independent MD replicates (left to right). The four views are the raw MD (ViViT) protein embedding (dark blue) and the raw ESM-2 sequence embedding (dark red)—both before contrastive learning—and the MD-projected embedding (light blue) and the DPLM-projected sequence embedding (light red)—both after contrastive learning; all four views are embedded together in a single cosine-UMAP per replicate. Before projection, the raw MD and raw ESM-2 clouds occupy well-separated regions, reflecting their distinct origins. After projection, the MD-projected and DPLM-projected clouds move together into neighbouring regions, showing that the contrastive objective reshapes the sequence representation to align with molecular-dynamics structure. The clouds converge into adjacent regions rather than collapsing onto one another because each protein contributes several MD replicates whose embeddings differ. This visualization corresponds to the cross-modal distance quantified in main-text Fig. 2d.

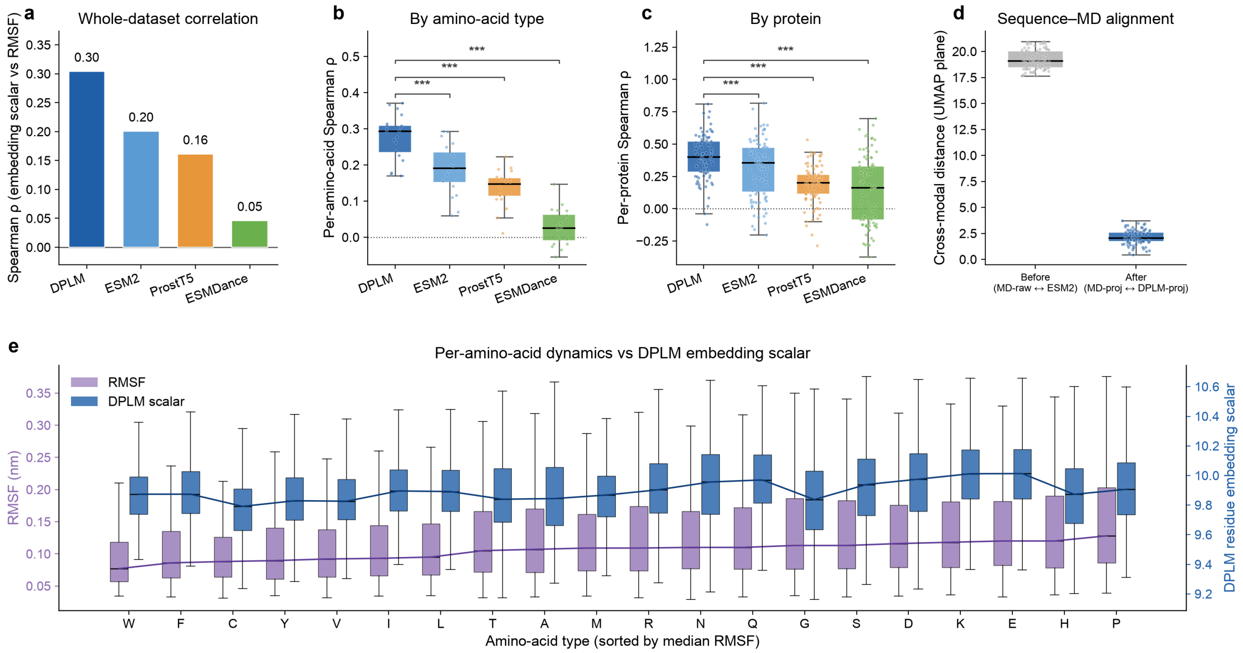


**Fig S4. Residue- and protein-level dynamics analysis for MD replicate 1.** Same analysis and layout as main-text Fig. 2, recomputed using the first of the three independent ATLAS molecular-dynamics replicates. (a) Whole-dataset Spearman ρ between the per-residue embedding scalar (the L2 norm of each model's per-residue representation) and RMSF, pooled over all residues of the test proteins, for DPLM, ESM2, ProstT5 and ESMDance. (b) The same correlation computed within each amino-acid type and summarized across the 20 types. (c) The correlation computed within each test protein and summarized across proteins; boxes show the median and interquartile range, points individual amino acids (b) or proteins (c). (d) Cross-modal distance between paired sequence and MD embeddings in a joint cosine-UMAP plane, before and after the contrastive projection. (e) Per-amino-acid distributions of RMSF (left axis) and the DPLM residue embedding scalar (right axis), amino acids ordered by increasing median RMSF, solid lines tracing the per-type medians (across-type Spearman ρ = 0.60, Pearson r = 0.56). DPLM shows the strongest embedding-scalar–flexibility coupling across all panels, reproducing the main-text result on this replicate.

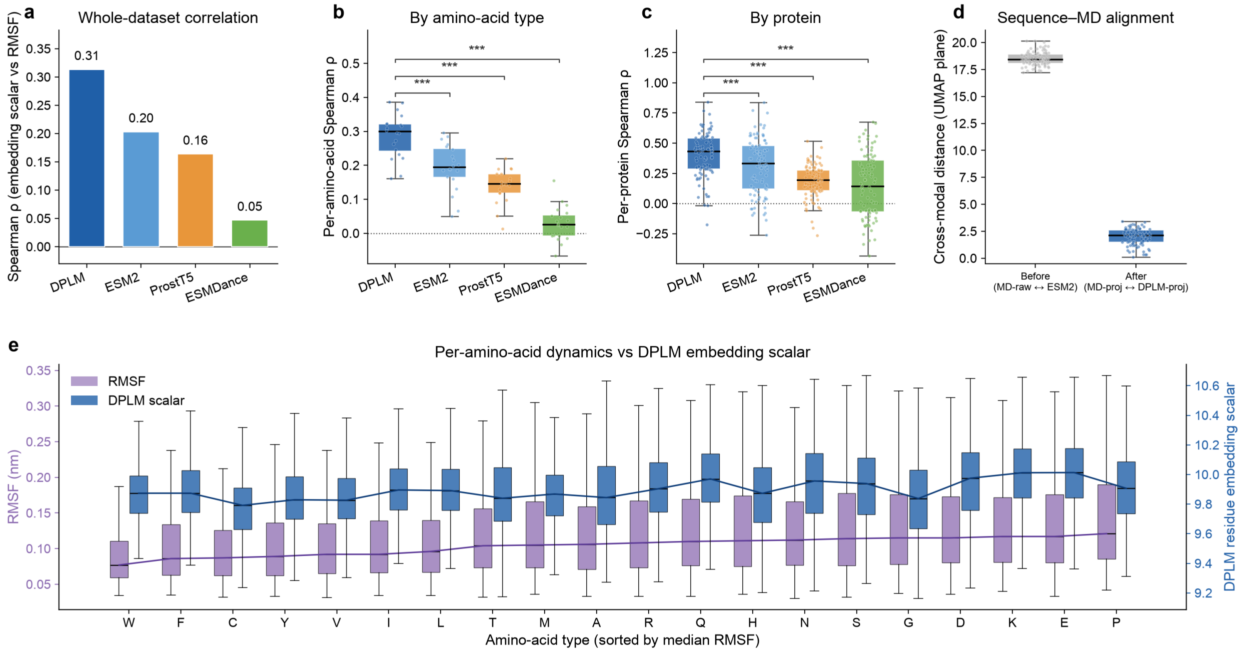


**Fig S5. Residue- and protein-level dynamics analysis for MD replicate 2.** Same analysis and layout as main-text Fig. 2, recomputed using the second of the three independent ATLAS molecular-dynamics replicates. (a) Whole-dataset Spearman ρ between the per-residue embedding scalar (the L2 norm of each model's per-residue representation) and RMSF, pooled over all residues of the test proteins, for DPLM, ESM2, ProstT5 and ESMDance. (b) The same correlation computed within each amino-acid type and summarized across the 20 types. (c) The correlation computed within each test protein and summarized across proteins; boxes show the median and interquartile range, points individual amino acids (b) or proteins (c). (d) Cross-modal distance between paired sequence and MD embeddings in a joint cosine-UMAP plane, before and after the contrastive projection. (e) Per-amino-acid distributions of RMSF (left axis) and the DPLM residue embedding scalar (right axis), amino acids ordered by increasing median RMSF, solid lines tracing the per-type medians (across-type Spearman $\rho = 0.63$, Pearson $r = 0.60$). DPLM shows the strongest embedding-scalar–flexibility coupling across all panels, reproducing the main-text result on this replicate.

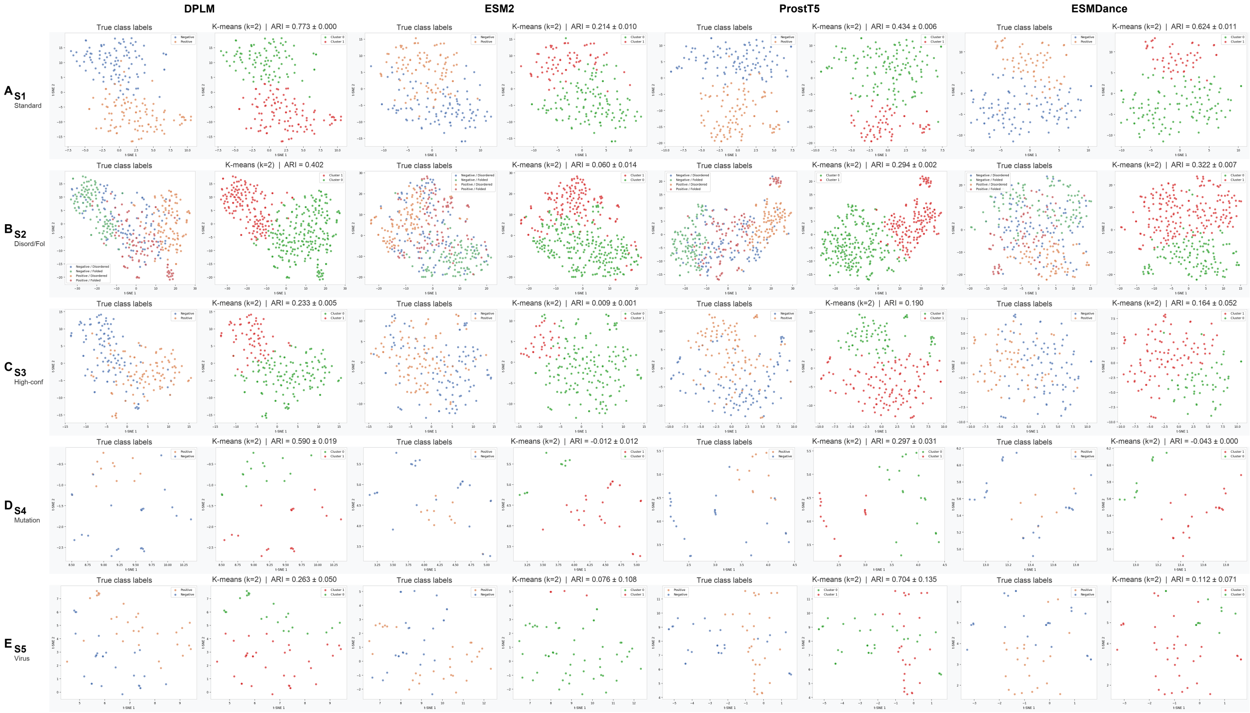


**Fig S6.Unsupervised recovery of phase-separation labels across embedding models.** t-SNE projections of protein embeddings from DPLM, ESM2, ProstT5, and ESMDance for the five phase-separation datasets (rows A–E: Standard, Disordered/Folded, High-confidence, Mutation, Virus). For each model, the left panel is colored by experimental LLPS label and the right panel by unsupervised k-means clusters (k = 2), with the adjusted Rand index (ARI) against the true labels shown above each k-means panel. DPLM yields the highest ARI on four of the five datasets (the structure-aware ProstT5 leads on the viral set S5), indicating that phase-separation-relevant structure is more strongly organized in the dynamics-aware representation than in the sequence-only (ESM-2), structure-aware (ProstT5), or alternative dynamics-informed (ESMDance) embeddings.

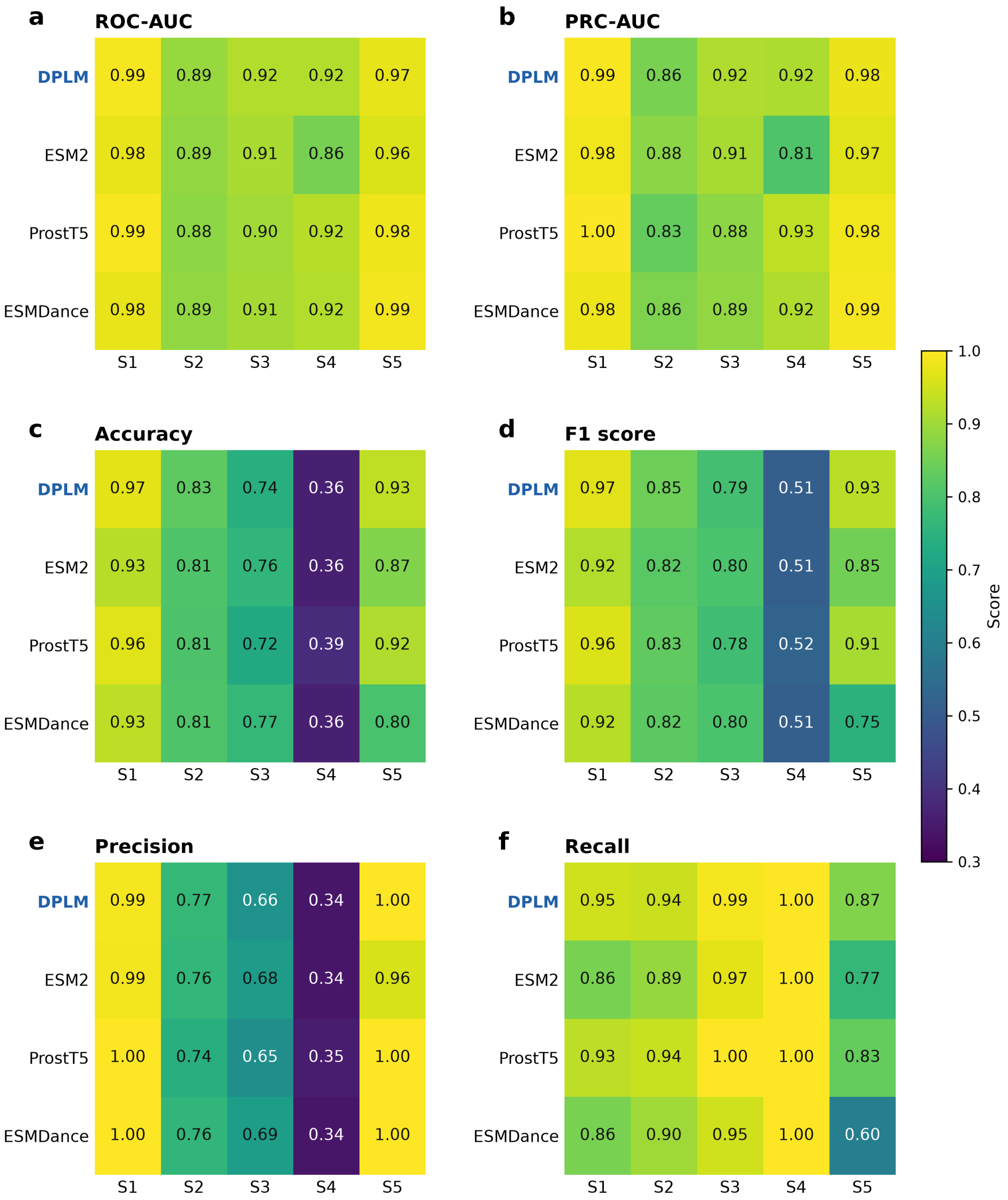


**Fig S7. Per-dataset supervised classification metrics for protein language model representations on phase-separation benchmarks.** ROC-AUC (a), PRC-AUC (b), accuracy (c), F1 score (d), precision (e) and recall (f) for a shared classifier head trained on the frozen embeddings of each protein language model (DPLM, ESM-2, ProstT5, ESMDance), evaluated per dataset across the five phase-separation benchmarks S1–S5 [Name Year]. Cell colour and printed value give the metric on each dataset; these are the per-dataset scores summarized as S1–S5 means in Fig. 3c. On the ranking metrics (ROC-AUC, PRC-AUC) the four models are near-indistinguishable across all datasets. On the threshold metrics (accuracy, F1, precision, recall) DPLM is highest or tied-highest on most datasets, with the strongest recall overall. All models degrade on S4, the smallest and most class-imbalanced benchmark (N = 36; 12 positive / 24 negative), where accuracy, F1 and precision fall for every model while recall stays saturated, a signature of majority-class-biased predictions on a small test set rather than a model-specific failure.

**Table S1. The filtered datasets statistics**

| Benchmark | Unit | Full set | Filtered set | Removed |
|---|---|---|---|---|
| S669 | proteins | 93 | 84 | 9 |
| CryptoBench (test) | apo chains | 222 | 176 | 46 |
| Mega-scale | proteins | 412 | 387 | 25 |
| Designed protein | proteins | 146 | 146 | 0 |
| Viral protein | proteins | 18 | 10 | 8 |
| S1 | proteins | 200 | 177 | 23 |
| S2 | proteins | 400 | 377 | 23 |
| S3 | proteins | 200 | 184 | 16 |
| S4 | proteins | 36 | 36 | 0 |
| S5 | proteins | 60 | 45 | 15 |

Each dataset was filtered against Atlas training set to make sure any pair of proteins from the two sets doesn't have a protein sequence identity >50%.

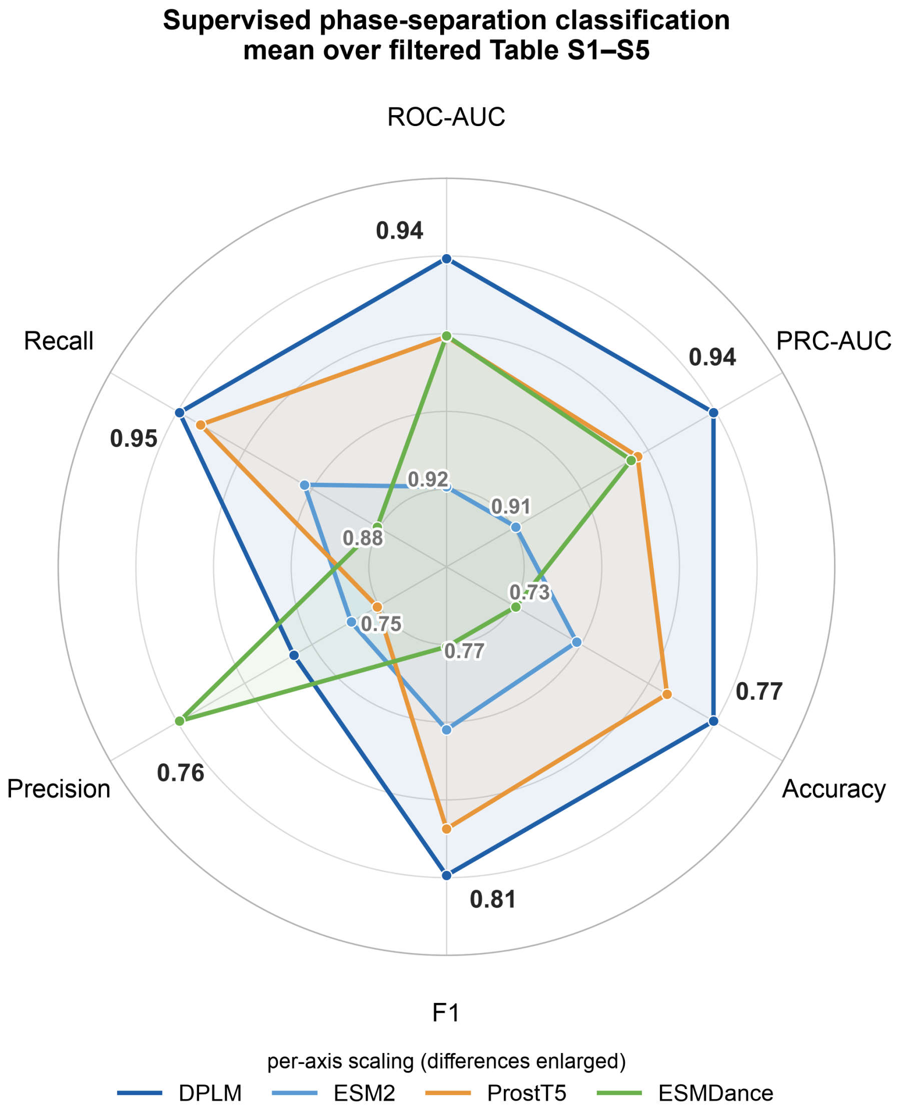


**Fig S8. Supervised classification with an identical classifier head on top of each language model.** Six metrics averaged over the filtered S1-S5 (max and min values for each metric is annotated in the radar plot).

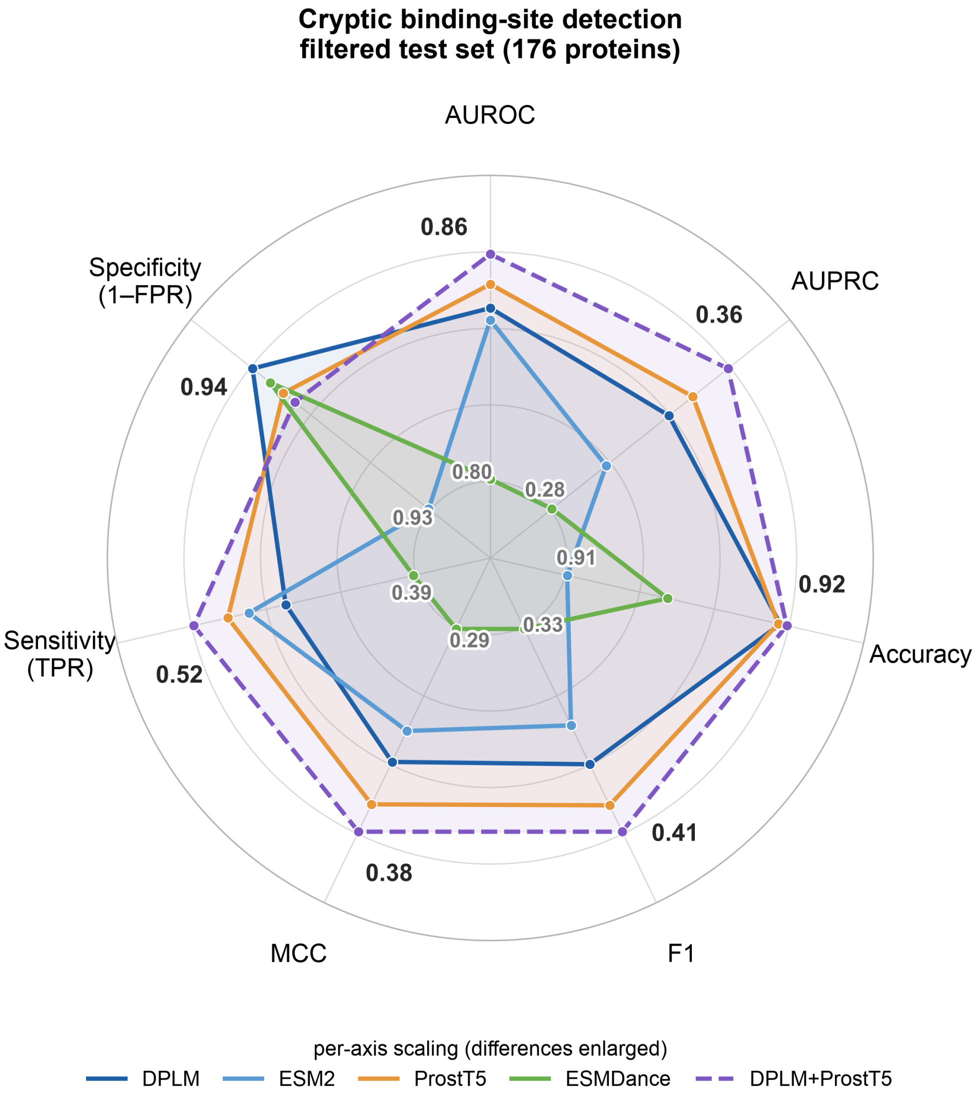


**Fig S9. Filtered CryptoBench cryptic-site detection scored per residue across seven metrics.** For four single encoders, DPLM, ESM2, ESMDance, and ProstT5, and one feature-fusion classifiers that concatenate DPLM embeddings with ProstT5 (single models solid, fusions dashed). Each axis is independently scaled to the observed range (per-axis min and max printed at the spoke ends) to make small differences visible.

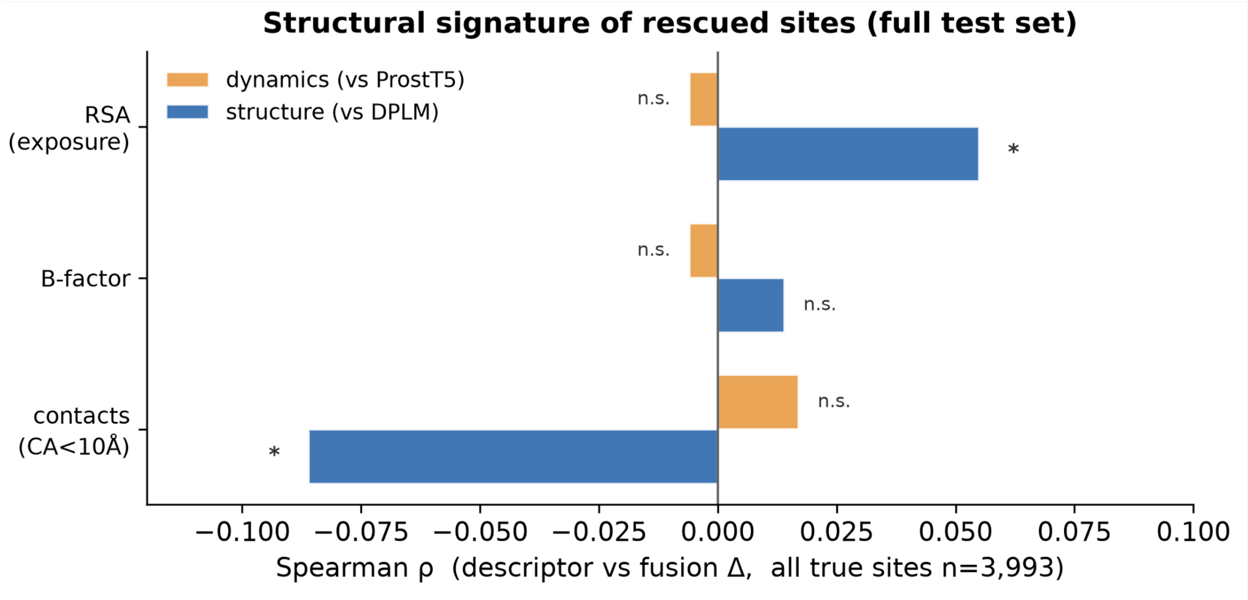


**Fig S10. Structural signature of the rescued residues over all 3,993 true sites.** Spearman ρ between each residue's fusion-score gain and three local structural descriptors.

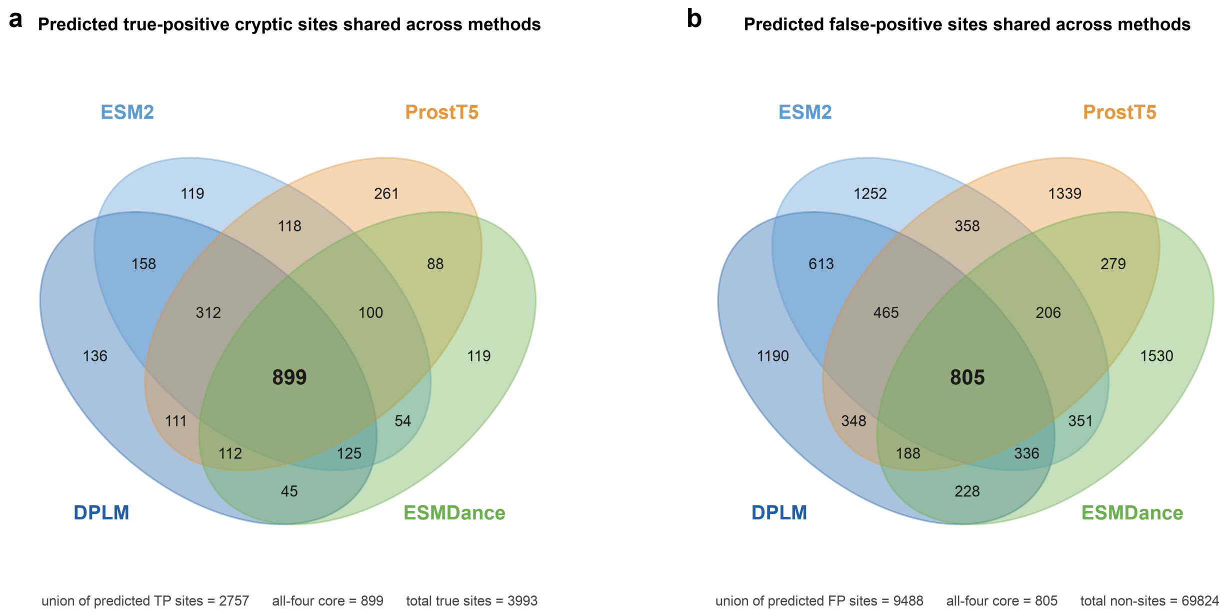


**Fig S11. The Venn diagram of overlap between individual PLM predicted cryptic site using the cryptobench dataset.**
**a.** true positive. **b.** false positive.

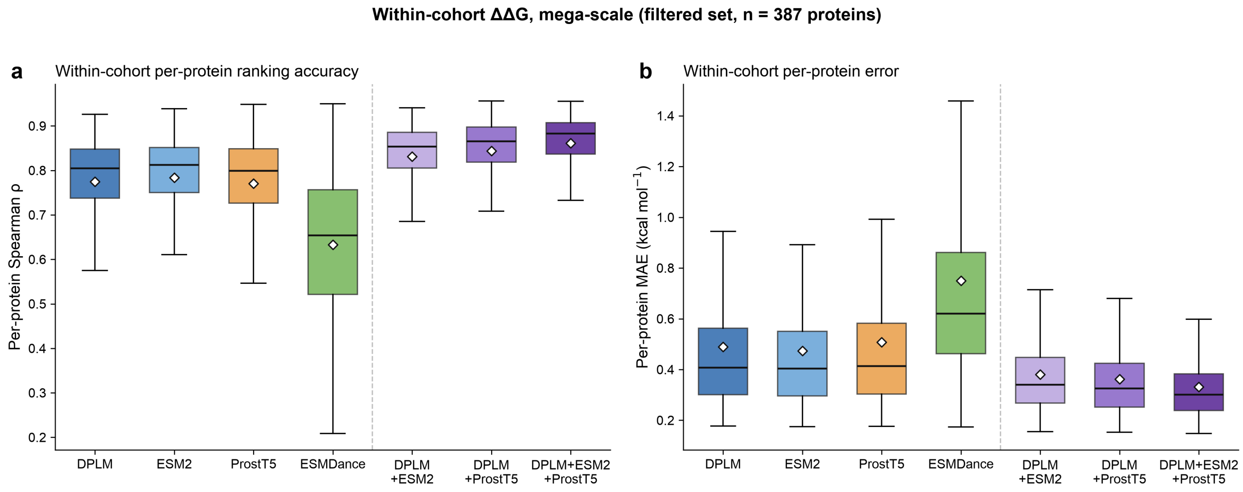


**Fig S12. Within-cohort ΔΔG prediction on the filtered mega-scale folding-stability dataset (387 proteins), scored per protein**. a. Spearman ρ (ranking; higher is better) and b. mean absolute error (kcal mol⁻¹; lower is better), for the four single models and three feature-fusion classifiers (DPLM+ESM2, DPLM+ProstT5, DPLM+ESM2+ProstT5).

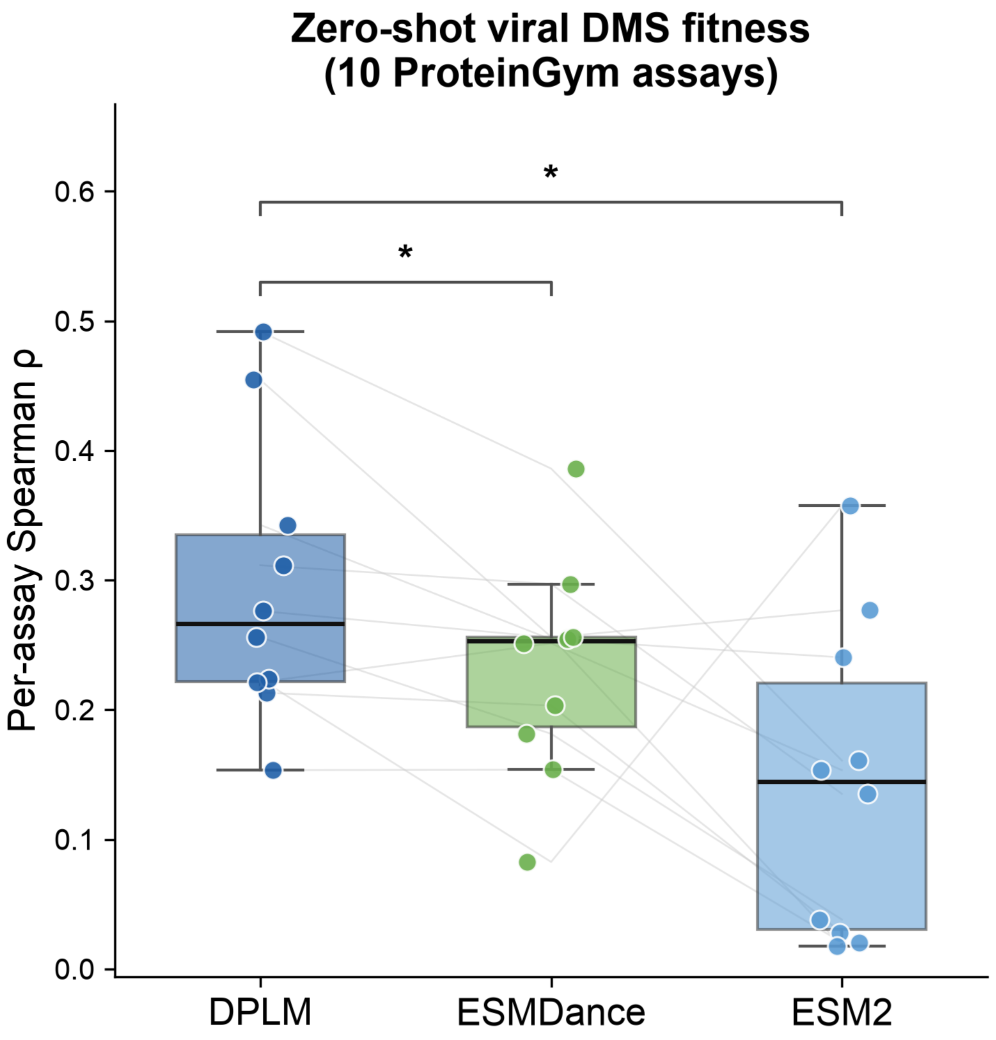


**Fig S13. Zero-shot variant-effect prediction on the filtered viral deep-mutational-scanning assays (10 assays) from ProteinGym, scored per assay by Spearman ρ between model score and measured fitness.** Each point is one assay; thin lines connect the same assay across models.

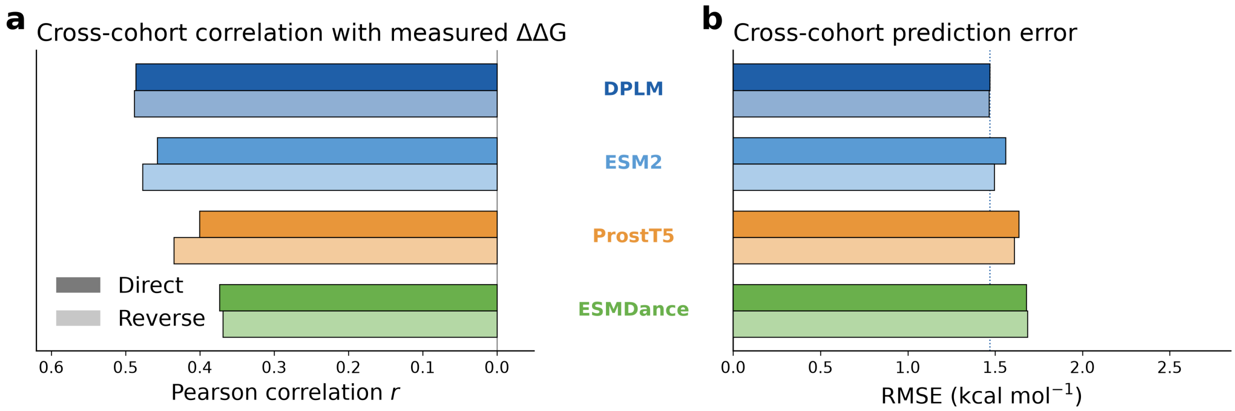


**Fig S14. Stability change prediction performance on the filtered S669 dataset. a.** Pearson correlation r and **b.** RMSE (kcal mol$^{-1}$) for the direct (darker color) and reverse (lighter color) mutation sets; the dotted line in (**b**) marks DPLM's direct RMSE.
